# Challenges and Opportunities in Single-Sample Network Modeling

**DOI:** 10.64898/2026.02.27.708608

**Authors:** Marieke Kuijjer, Margherita De Marzio, Kimberly Glass

## Abstract

Analysis of biological networks can provide unprecedented insights into the mechanisms underlying disease. Although many methods have been developed to estimate biological networks, these approaches typically use multiple experimental samples to estimate a single ‘aggregate’ network, which fails to capture population-level heterogeneity. Recently, several methods have been developed that overcome this limitation by inferring networks for individual samples, i.e. single-sample networks. However, each approach for inferring single-sample networks has been formulated differently, making it challenging to compare them. To address this issue, we re-cast the mathematics of several single-sample network methods using common variables. We then systematically explore the parameters, caveats, and underlying assumptions made by each method and examine how these differences impact single-sample network prediction. Our analyses point to a critical trade-off that occurs when trying to simultaneously predict network edges that are both shared across samples as well as edges that are specific to a given sample. For example, the mathematics of both SWEET and BONOBO includes a scale factor that drives the weights of edges in the predicted single-sample networks toward a background network. The result is that, although networks predicted by these methods tend to have the highest accuracy, this often comes at the cost of very low specificity, an important caveat since the primary goal of sample-specific network modeling is to obtain networks that are *specific* to each input sample. In contrast, SSN estimates the most specific but least accurate networks, while LIONESS straddles these domains, with an accuracy almost as high as SWEET and BONOBO and a specificity almost as high as SSN. Overall, our analyses highlight some of the broader challenges in this emerging field. However, they also point to important methodological synergies, providing an opportunity to create a common framework that can be used to improve single-sample network inference.

## 2 Introduction

Network modeling approaches provide an important framework for understanding the mechanisms that drive biological systems. Many methods have been developed to infer biological networks, often by leveraging data from multiple samples to construct a single representative model [1, 2, 3, 4]. Analyzing a single or handful of networks representing different biological states can provide important insights into health and disease [5, 6]. However, this type of approach is limited in its ability to provide insight into how networks vary across a population [7]. Some approaches overcome this limitation by layering omics data onto a known network structure (such as a reference protein-protein interaction or gene regulatory network); these methods then calculate sample-specific scores for nodes [8] or modify nodes/edges in the reference network based on data from individual omics samples [9, 10, 11]. In this paper, we focus on a distinct set of algorithms that can be used to directly infer values (or weights) for all possible edges in a single-sample network. Although most often applied to gene expression data [12], these types of single-sample network methods have also been used to analyze other types of omics data, such as metabolomics [13][14], microbiome [15, 16, 17, 18], and genotype [19] data.

The first methods proposed to directly infer sample-specific networks included LIONESS (Linear Interpolation to Obtain Network Estimates for Single Samples) [20] and SSN (Sample Specific Network) [21]. These two methods were developed independently and posted on pre-print servers within one year of each other [22, 23]. Both LIONESS and SSN can be thought of as mathematical formulas for estimating a sample-specific network. The LIONESS equation was derived in a method-agnostic manner and models a sample-specific network that includes both shared information (i.e. edges that are common across the input population) and sample-specific information (i.e. edges that are specific to an individual sample). This means that the edge values predicted by LIONESS depend on an accompanying ‘aggregate’ network inference algorithm. In their paper, Kuijjer et al. [20] applied LIONESS to networks derived using Pearson correlation, Mutual Information (MI), PANDA [4], and CLR [24], four methods that return a single network based on information from all samples in an input dataset. Others have applied LIONESS to networks constructed using many different algorithms, including partial correlation [18], MI/ARACNe [25], PANDA [26, 27, 28], principal components [17], and rMAGMA [15, 16].

In contrast, SSN was developed specifically to infer sample-specific *correlation* networks. SSN leverages a statistical framework (Z-score) to capture how much the addition of a single sample to a dataset perturbs an original ‘reference network’. Thus, the edge values predicted by SSN look like Z-scores. In contrast to LIONESS, SSN infers only sample-specific network information (rather than both shared and sample-specific information). Another related approach, SWEET (sample-specific weighted correlation network) [29], uses a modified version of the LIONESS equation that aims to account for subpopulation structure. In their paper, Chen et al. only apply SWEET to networks derived from Pearson correlation. Another recent method, BONOBO (Bayesian Optimized Networks Obtained By Assimilating Omics) [30] uses a Bayesian model to derive sample-specific correlation matrices. BONOBO is the only approach whose predicted sample-specific networks are designed to have statistical properties that resemble Pearson correlation. Together, SSN, SWEET, and BONOBO represent three methods specifically developed or presented with the goal of inferring sample-specific correlation networks. In this paper, we explore these three methods in depth, and compare their predictions to those made by LIONESS when applied to Pearson correlation (referred to as LIONESS::PCC).

Several single-sample network methods have also been developed that can capture more complex relationships. For example, as noted above, in their paper, Kuijjer et al. [20] apply LIONESS to MI networks. CSN (cell specific network) [31], initially developed for application to single-cell gene expression data, also leverages an approach reminiscent of MI estimation, binning expression values to estimate probability distributions for each gene; these distributions are used to define a cell-specific normalized joint probability for gene pairs. The mathematics of CSN has also been applied to infer sample-specific networks using bulk expression data [32]. In this context, each ‘cell’ in CSN is a ‘sample’ in the bulk dataset. In this paper, we explore both of these ‘non-linear approaches’: LIONESS applied MI networks (referred to as LIONESS::MI) and CSN (in the context of bulk data).

Despite their common goal, the mathematics of these methods was formulated in an inconsistent manner, with different variable names used to represent similar concepts. This has made it challenging to systematically evaluate the similarities, differences, and strengths of these single-sample network methods. Several benchmarking studies have recently compared the performance of these methods in various real-world settings [32, 14, 33]. Although the results of these studies fill a critical gap by providing an independent characterization of the methods’ predictions, they do not provide an intuition as to why one method may perform better (or worse) in a particular context.

In this paper, we explore the *mathematical framework* for five single-sample network methods: LIONESS, SSN, SWEET, BONOBO, and CSN. By re-casting the equations that define these methods using common variables, we are able to more clearly delineate their similarities and characterize their key differences. This also allows us to examine the parameters, caveats, and underlying assumptions made by each method and determine how these may impact single-sample network prediction. In contrast to previous benchmarking studies, which have focused on method accuracy or predictive capacity, the primary goal of this study is to identify common mathematical themes across single-sample network approaches and to provide key insights into how different method parameters and data features might *impact* method predictions and perceived performance. Our goal is to help others develop an intuition of how single-sample network methods work by providing a consistent mathematical formulation of the methods, an understanding of the methods’ parameters, and a characterization of the impact of those parameters on single-sample network prediction.

## 3 Similarities (and Differences) of Single-Sample Network Methods

Many single-sample network methods have either been explicitly derived in the context of Pearson correlation (SSN, BONOBO) or can be applied to Pearson correlation networks (LIONESS::PCC, SWEET). In Figure 1 we show the equations that define a sample-specific network for each of these methods. These equations have been simplified and recast to use the same variables and conceptualization framework. For more information, see Supplemental File 1.

**Figure 1.**
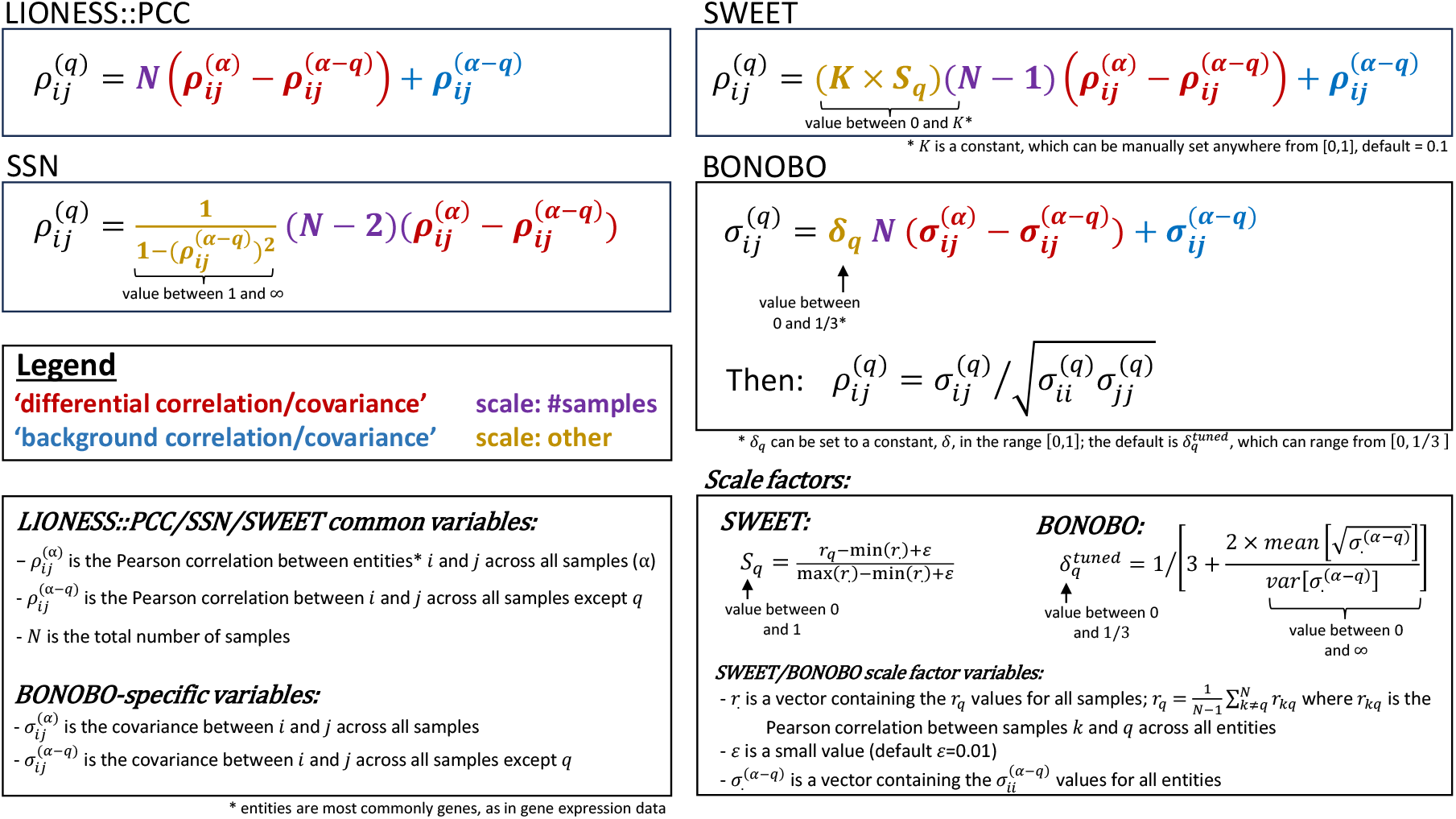
Mathematical formulation of four single-sample network methods based on Pearson correlation.

Inspection of these equations reveals that these four methods have a similar mathematical formula that includes up to four components: (1) a differential correlation (or covariance) term (red), (2) a scale factor representing the number of samples (purple), (3) an additional scale factor (gold), and (4) a background correlation (or covariance) term (blue). These components are also organized similarly: all methods multiply the differential correlation / covariance term by a scale factor related to the number of samples, all methods except for LIONESS::PCC further multiply this term by an additional scale factor, and all methods except for SSN add in a background correlation / covariance term. In contrast to the other methods, BONOBO is explicitly designed to generate edge weights that resemble a Pearson correlation. It does this by first estimating an unbounded sample-specific covariance matrix; BONOBO then transforms this sample-specific covariance matrix into a sample-specific correlation matrix, which is bounded between −1 and 1.

The most significant difference in these equations is in the formula for the additional scale factor that multiplies the differential correlation / covariance term in SSN, SWEET, and BONOBO but not LIONESS::PCC. The motivation behind the additional scale factor is different for each method: for SSN the scale factor is a result of the statistical derivation, for SWEET the scale factor captures population substructure, and for BONOBO the scale factor represents the relative sample-specific (percent) contribution (see Supplemental File 1). Consequently, these additional scale factors are formulated very differently. For SSN, this factor scales based on the background correlation, and can vary in value between 1 and ∞. For SWEET, the scale factor is composed of two variables: (1) a ‘balance parameter’ *K*, and (2) a sample-specific weight *S*_*q*_. Chen et al. set *K* = 0.1 in their study while *S*_*q*_ represents a sample-specific similarity score that varies between 0 and 1. Thus, the combined scale factor for SWEET, *S*_*q*_ × *K*, is bounded between 0 and *K* (by default, 0 and 0.1). Similarly, in BONOBO, the scale factor, *δ*_*q*_, is a sample-specific weight. By default, BONOBO sets 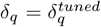, which can take a value between 0 and 1*/*3, although Saha et al. [30] state that *δ*_*q*_ can alternately be set equal to any constant value between 0 and 1. The *K, S*_*q*_, and *δ*_*q*_ variables are examined in detail in Section 5.

It is important to note that, in addition to differences in their mathematics, there are also important differences in how single-sample methods have been conceptualized. For example, LIONESS, BONOBO, and CSN were all originally conceptualized as ‘leave-one-out’ approaches, meaning that they first compute an ‘aggregate network’ based on all samples and then remove one sample to compute a ‘background network’; the aggregate and background networks are then used to estimate the network for the left-out (removed) sample. In contrast, SSN was originally conceptualized as an ‘add-one-in’ approach. In this case, a set of samples is used to calculate a ‘reference network’. A single sample is then added to this reference set and the combined samples are used to estimate a ‘perturbed network’; the reference and perturbed networks are then used to estimate the network for the added (non-reference) sample. SWEET takes a third approach. In SWEET, all samples from the input dataset are used to compute a reference network. Each input sample is then iteratively re-added to the data (i.e. a sample-of-interest is duplicated in the expression data) and the combined samples are used to estimate a perturbed correlation network; the single-sample network is then estimated for the re-added sample. As has been pointed out by others [32], the ‘leave-one-out’ approach is the most versatile since often true reference populations are limited in size and defining the correct reference for an analysis can be tricky. Therefore, for consistency, we re-implemented all of these methods based on the ‘leave-one-out’ conceptualization (see Supplemental File 1). However, in Supplemental File 2 we explore the impact of implementing these methods using an ‘add-one-in’ approach; we also demonstrate that the results presented in this manuscript for SWEET are essentially identical when using the standard ‘leave-one-out’ approach compared to the sample duplication approach used in SWEET’s original conceptualization.

## 4 Comparing the Predicted Networks

To evaluate how differences in the formulation of single-sample network methods impact the prediction of sample-specific network edge weights, we derived a toy gene expression dataset. As illustrated in Supplemental Figure 1A, this toy data is composed of six genes and 600 samples within which there are three distinct ‘known’ correlation networks: (1) a fully connected network (Sample Set 1: all genes correlated; 250 samples), (2) a network with two cliques (Sample Set 2: two sets of genes correlated with anti-correlation between the sets; 250 samples), and (3) a fully disconnected network (Sample Set 3: no correlation between any genes; 100 samples). This data gives rise to two primary edge types, as defined by their pairwise gene correlation patterns across all samples: (1) a linear edge (plus noise from Sample Set 3) and (2) a non-linear edge (plus noise from Sample Set 3). Two exemplar edges, one for each type, are shown in Supplemental Figure 1B.

We applied six single-sample network methods to these toy data and visualized the edge weights predicted by each method for all samples and two exemplar edges. This included four ‘linear’ methods (LIONESS::PCC, SSN, SWEET, BONOBO) and two ‘non-linear’ methods (LIONESS::MI, CSN). In our visualization, we used both an ‘automatic’ color range (Figure 2A), which is different for each plot and ranges from the minimum to the maximum value predicted across all samples for that edge and method, and a ‘consistent’ color range (Figure 2B), which always ranges from −2 to 2. The ‘automatic’ color range allows us to observe trends within each method and edge type, while the ‘consistent’ color range allows us to visually compare the predicted edge weights. Direct pairwise comparisons of predicted edge weights between methods and edge types are provided in Supplemental Figure 2. Edge weights for all samples, edges, and methods are visualized using the ‘consistent’ color range in Figure 2C.

**Figure 2.**
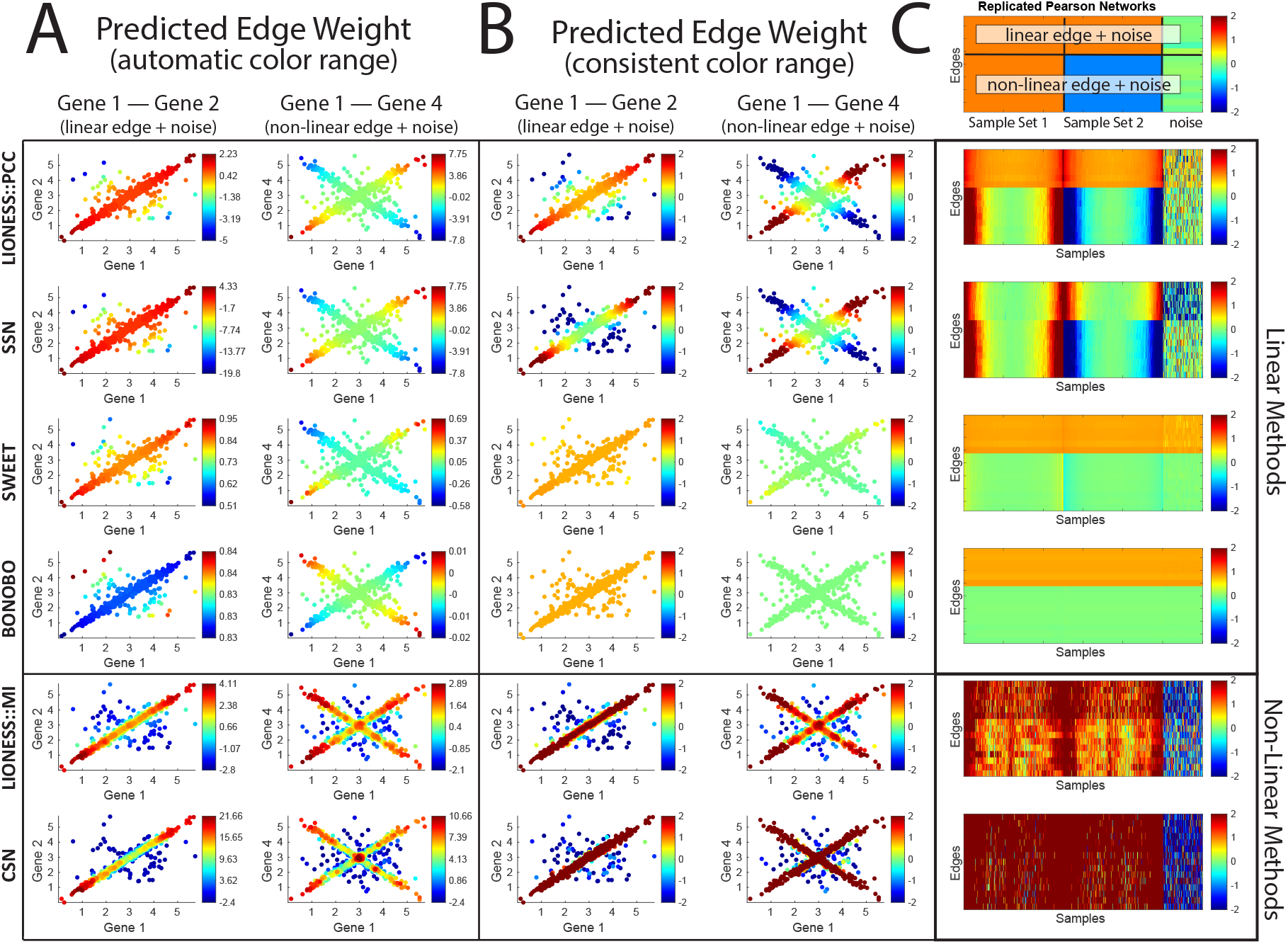
The edge weights predicted for each of the samples that make up two exemplar edges when applying six different single-sample network methods to the toy gene expression data. The edges visualized include: (1) Gene 1 – Gene 2, which have an overall linear relationship across all samples, including some ‘noise’ from samples in sample set 3; and (2) Gene 1 – Gene 4, which have an overall non-linear relationship across the samples, including some ‘noise’ from samples in sample set 3. (A-B) Each point is a sample and the predicted edge weights for each method are visualized using either (A) an automatic color range that differs for each edge and method and ranges from the minimum to the maximum value predicted across all samples for that edge and method, and (B) a consistent color range that is the same for each edge and method; samples with edge values outside the range are given the same color, either blue (weight*<* − 2) or red (weight*>* 2). (C) A heat map showing the predicted weight for all edges across all samples when applying each of the six different single-sample network methods. Within sample set 1 and sample set 2, samples are in ascending order based on the expression of genes 1-3 (see Methods). At the top of (C) is a heat map showing the expected ‘reference’ value of these edges in each of the three sample sets. See also Supplemental Figure 1.

In these plots, we observe many similarities as well as important differences in the predictions made by the single-sample network methods. For example, the edge weights predicted by LIONESS::PCC and SSN are perfectly correlated for both exemplar edges (Figure 2, top two rows; Supplemental Figure 2). However, the values predicted by these methods differ for the linear edge. In particular, for LIONESS::PCC, samples along the *x* = *y* diagonal have values close to one or slightly above, while for SSN these samples have a value close to zero or slightly below. The overall range of values for the linear edge is also larger for SSN compared to LIONESS::PCC. These differences do not occur for the non-linear edge. These results can be understood by examining the mathematics underlying these two methods (see Figure 1). For the linear edge, the correlation of the genes that make up the edge is close to one, thus the scale factor for SSN becomes large, increasing the impact of the ‘differential correlation’ term in SSN and, consequently, the range of predicted values. In addition, the ‘background correlation’ term of LIONESS::PCC essentially centers its predicted values around one (instead of zero as is the case for SSN). On the other hand, for the non-linear edge, the correlation of the genes that make up the edge is close to zero, so the scale factor of SSN approaches one, the ‘background correlation’ term of LIONESS::PCC approaches zero, and the SSN and LIONESS::PCC equations become essentially equivalent.

We next observe that the edge weights predicted by SWEET have a similar, but non-identical, trend to those predicted by LIONESS::PCC and SSN. However, the range of edge weights predicted by SWEET compared to LIONESS::PCC and SSN is much smaller (Figure 2, third row). This, once again, can be understood by examining the mathematics underlying these methods. SWEET primarily differs from LIONESS::PCC and SSN in terms of its scale factor, *K* × *S*_*q*_. This scale factor is bounded between 0 and *K*, whose default value is 0.1. Thus, the range of values multiplying the ‘differential correlation’ term in SWEET is by default at least 10 times smaller than that of LIONESS::PCC and SSN. This causes the edge weights predicted by SWEET to approach the ‘background correlation’ value, which is close to one for the linear edge and close to zero for the non-linear edge.

We also observe that edge weights predicted by BONOBO are in a similar *range* as those of SWEET; all samples have an edge weight close to one for the linear edge and close to zero for the non-linear edge (Figure 2B-C, fourth row). However, strangely, the edge weights predicted by BONOBO are negatively correlated with those predicted by every other single-sample network approach (Figure 2A; Supplemental Figure 2). Counterintuitively, for the linear edge BONOBO predicts that samples closest to the *x* = *y* diagonal have the lowest weights, and for the non-linear edge BONOBO predicts that samples at the extreme of the *x* = *y* line have the lowest weights while those at the extreme of the *x* = − *y* line have the highest weights. This is the opposite pattern as what we observe for LIONESS::PCC, SSN, and SWEET. We explore the reason for this observation in the following section.

Finally, we observe that the edge weights predicted by the two non-linear approaches, LIONESS::MI and CSN, are highly correlated, although the edge weights predicted by CSN have a higher range across the samples compared to LIONESS::MI (Figure 2, fifth and sixth row; Supplemental Figure 2). For the non-linear edge, LIONESS::MI and CSN generally give high weight to the samples that make up the non-linear pattern (sample set 1 with sample set 2) and low weights to ‘noisy’ samples that are not part of this pattern (those in sample set 3). This is in contrast to LIONESS::PCC and SSN, which, for the non-linear edge, give highly positive edge weights to samples along the extremes of the *x* = *y* diagonal, highly negative edge weights to samples along the extremes of the *x* = −*y* diagonal, and edge weights close to zero near the origin.

## 5 The Role of Additional Parameters

Both SWEET and BONOBO include a scale factor that is not derived from any of the common variables shared across the methods (Figure 1). We systematically evaluated these scale factors to determine how they impact the predicted edge weights.

We first examined *δ*_*q*_ in BONOBO, which by default is set equal to 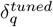 (Figure 1). The value of 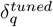 is derived using information from all genes in the input data set, captured in a vector *σ*^(*α−q*)^ whose elements contain the variance of each gene’s expression across all samples except *q*. Although Saha et al. [30] state that *δ*_*q*_ can be manually set equal to any constant value between 0 and 1, 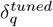 has a theoretical limit of 1*/*3. This is because the value of the second term in the denominator of the equation defining 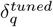 (Figure 1) ranges between zero and ∞. To generate the toy data we used a normal random distribution function (see Methods). The (unintended) consequence was that every gene in these data had approximately the same variance across samples. In other words, var[*σ*^(*α−q*)^] was always very close to zero, causing 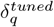 to be very close to zero, and leading to the edge weights predicted by BONOBO being dominated by the ‘background covariance’ term.

With this in mind, we once again applied BONOBO to the toy data, but set *δ*_*q*_ = *δ*, a constant value across all samples. We observe that varying *δ* changes the range and distribution of the edge weights predicted by BONOBO. However, these edge weights are always perfectly non-linearly correlated with those of LIONESS::PCC (Figure 3A-B). As explained above, BONOBO generates edge weights that are bounded between −1 and 1. This is evident in the plots shown here and leads to the observed non-linear relationship between the edge weights predicted by BONOBO and LIONESS::PCC. We note that since LIONESS::PCC and SSN are perfectly correlated (see Supplemental Figure 2) these trends would be identical had we compared BONOBO to SSN.

**Figure 3.**
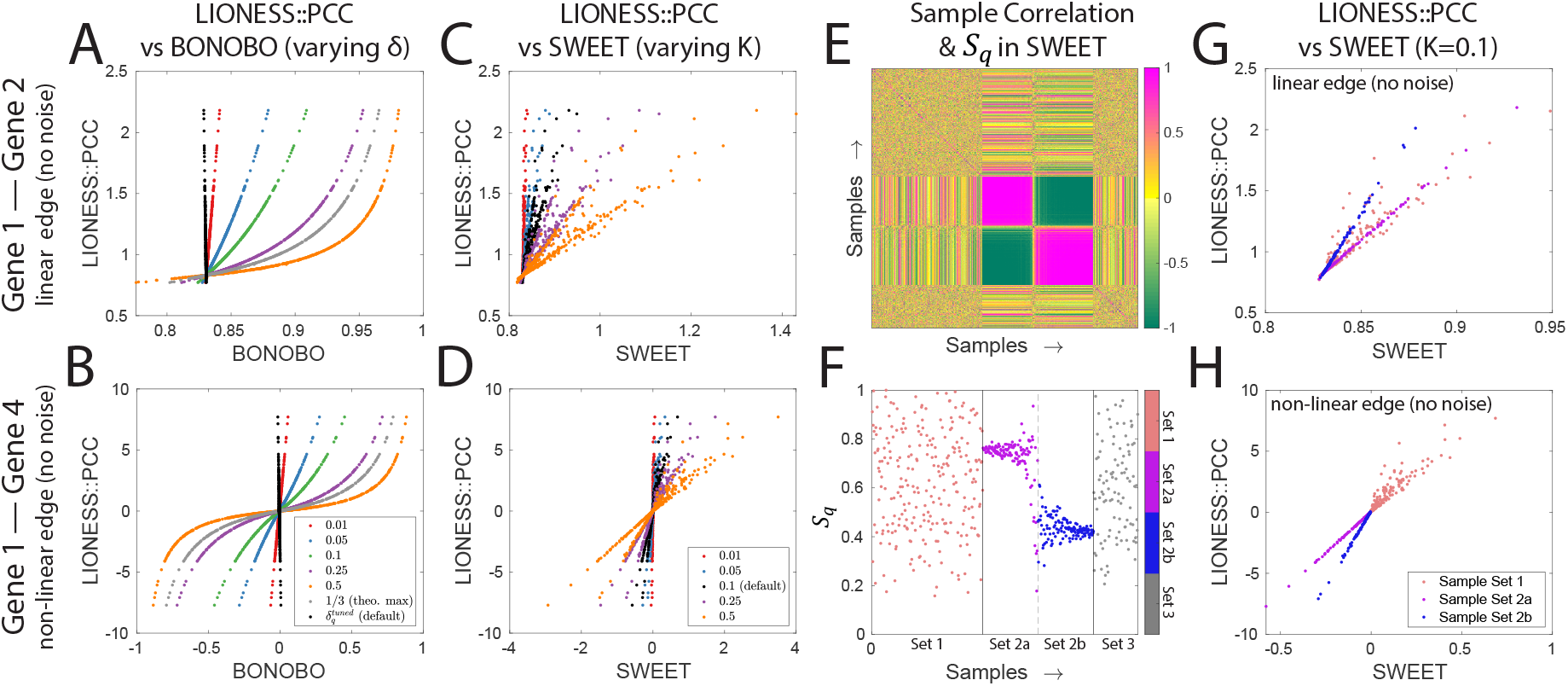
(A-B) A plot comparing the edge weights predicted by BONOBO for different values of *δ*_*q*_ versus LIONESS::PCC for both a (A) linear and (B) non-linear edge. (C-D) A plot comparing the edge weights predicted by SWEET for different values of *K* versus LIONESS::PCC for both a (C) linear and (D) non-linear edge. Note that since LIONESS::PCC and SSN are perfectly correlated (see Supplemental Figure 2), these relationships would be identical if comparing BONOBO and SWEET to SSN instead of to LIONESS::PCC. (E) The pairwise correlation between samples in the toy expression dataset. (F) The value of *S*_*q*_ in SWEET for each of the samples in the toy expression data. (G-H) A plot comparing the edge weights predicted by SWEET for *K* = 0.1 (default value) but colored based on sample set. Note that for all plots in this figure, we excluded samples from set 3 (noise) to better visualize the results. See also Supplemental Figure 2.

We next investigated the role of the scale factor in SWEET, which is composed of two variables: (1) a ‘balance parameter’ *K*, and (2) a sample-specific weight *S*_*q*_. In Chen et. al. [29] the authors state that *K* represents a percentage of samples. With this in mind, we once again applied SWEET to the toy data, but varied the value of *K*. We find that varying *K* in SWEET has a similar effect as varying *δ* in BONOBO, changing the range of the predicted edge weights and impacting the slope of association with LIONESS::PCC (3C-D). There is also a V-shape and X-shape pattern in these plots, which is visible across multiple values of *K*. We explore the reason for these patterns next.

SWEET also includes a variable, *S*_*q*_, which represents the normalized average similarity of a sample *q* with all other samples in the input data. Mathematically, *S*_*q*_ depends on the pairwise correlation between samples (*r*) (see equation in Figure 1). *S*_*q*_ = 1 for the sample with the highest average correlation with all other samples in the input data, while *S*_*q*_ → 0 for the sample with the lowest average correlation with all other samples in the input data, although the lower bound of *S*_*q*_ depends on the value of *ϵ*, which Chen et. al. set by default to 0.01 [30].

To understand the role of *S*_*q*_, we visualized the pairwise correlation between all samples (*r*) in the toy data (Figure 3E). We observe a striking pattern. Sample Set 2 separates into two distinct clusters, a grouping we did not intend when generating these data. The impact on *S*_*q*_ is pronounced (Figure 3F). Whereas in Sample Set 1 and Sample Set 3 the values of *S*_*q*_ are distributed fairly evenly between around 0.2 and 1, for samples in Sample Set 2 the values of *S*_*q*_ cluster around two values, ∼0.4 and ∼0.75. With this in mind, we again plotted the edge weights predicted by SWEET (using the default *K* = 0.1) against those predicted by LIONESS::PCC, but colored samples by their sample set (Figure 3G-H). We see that the V-shape and X-shape patterns we had previously observed are the consequence of the separation of Sample Set 2 into two groups. Following up on this observation, in the following section we systematically investigate how population substructure impacts single-sample method predictions.

## 6 The Impact of Data Substructure

Data substructures, including differences in expression, correlation, and sizes between subpopulations, are commonly encountered when investigating diseases, such as cancer, that have distinct subtypes. Therefore, we next evaluated single-sample network predictions using ‘real-world’ gene expression data containing two clearly defined subpopulations. We selected 500 gene expression samples from the Genotype Tissue Expression (GTEx) project [34], 250 samples each from two related but distinct tissues: esophagus mucosa and esophagus muscularis (Supplemental Figure 3A; see Methods). In these data, many genes are differentially expressed between tissues, but most edges (gene pairs) have a similar correlation value in both tissues (Supplemental Figure 3B).

Both SWEET and BONOBO, by default, include sample-specific scale factors (*S*_*q*_ and 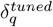) that are calculated using information from all genes in the input data (rather than just the two genes that make up a specific edge, see Figure 1). To characterize how these scale factors may be impacted by subpopulation size and data structure, we randomly sub-sampled the ‘real-world’ expression data described above such that each random sub-sample contained 100 random samples but varied in terms of the percentage of samples selected from each tissue, from 0% to 100% (examples are shown in Supplemental Figure 3C). We repeated this sub-sampling procedure 100 times for each percentage and calculated *S*_*q*_ and 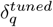 for each sample in the sub-sampled data. The results of this analysis show a clear impact on *S*_*q*_ (Figure 4A, Supplemental Figure 3C), with samples from the smaller population consistently having significantly lower *S*_*q*_ values than those from the larger population. In contrast, there are only minimal differences in the value of 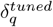 across samples in the smaller versus larger population (Figure 4B). Instead, this parameter is more sensitive to the overall composition of the data, with lower values of 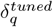 when a higher percentage of samples come from the same tissue.

**Figure 4.**
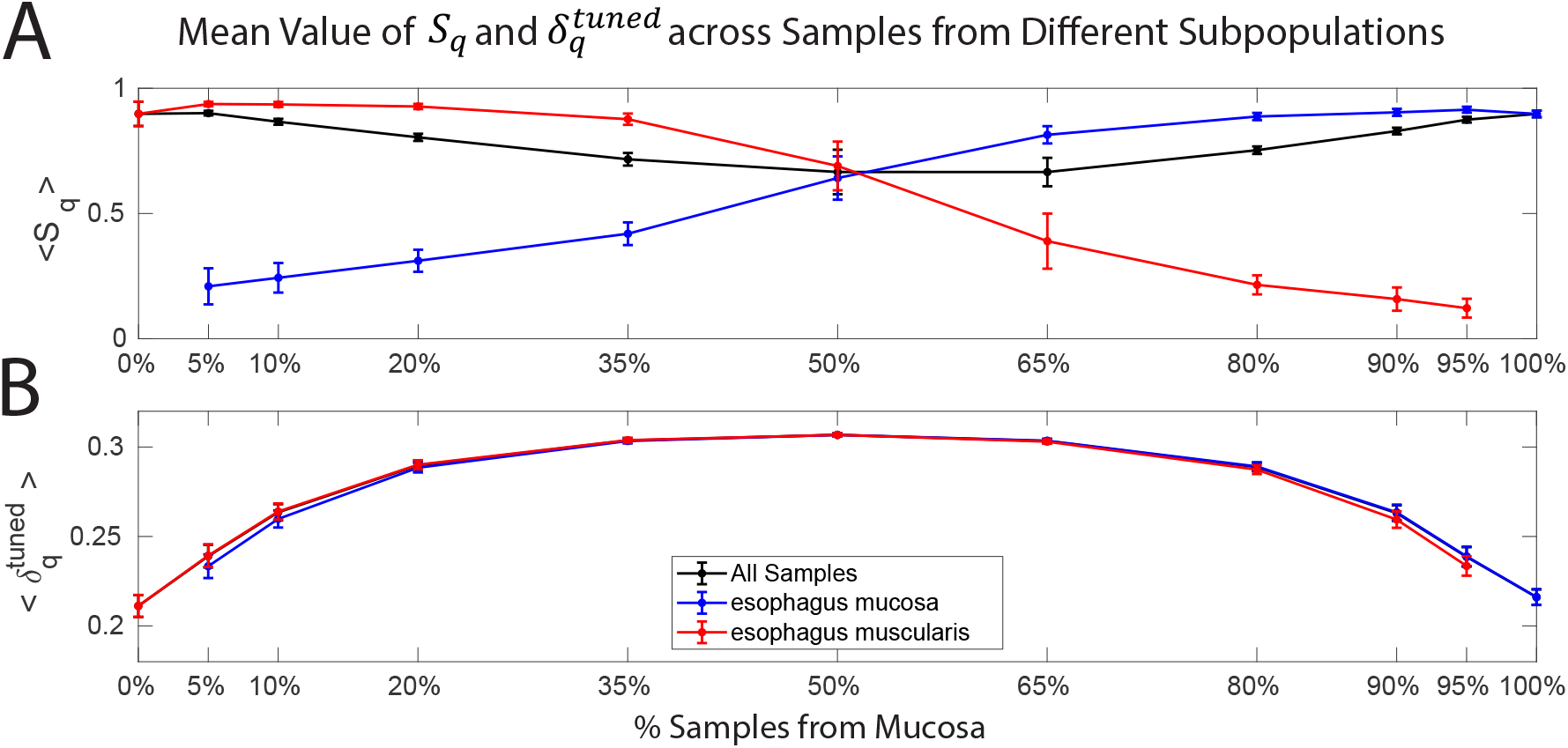
Evaluation of the *S*_*q*_ (SWEET) and *δ*_*q*_ (BONOBO) parameters in a ‘real-world’ gene expression dataset with two distinct subpopulations. The average value of (A) *S*_*q*_ and (B) 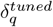 across samples from mucosa versus muscularis tissues as a function of the percentage of mucosa samples in the input data (random subset) used to infer the edge weights. The mean +/-standard deviation of these average values across 100 randomizations are shown with dots and error bars, respectively. See also Supplemental Figure 3.

We next evaluated how data substructure may impact predicted single-sample network edge weights. We focused on four exemplar edges (Figure 5A and Supplemental Figure 3B): (1) high correlation in both tissues and neither gene is differentially-expressed (similar to the linear edge in our toy data); (2) no correlation in either tissue and neither gene is differentially expressed; (3) high correlation in both tissues but in the opposite direction and neither gene is differentially-expressed (similar to the non-linear edge in our toy data); and (4) no correlation in either tissue but both genes are highly differentially expressed. These exemplar edges were selected to help disentangle the impact of differential correlation versus differential expression on the predicted edge weights. For each of these exemplar edges, we determined the edge weights predicted by each single-sample network method in each random sub-sample. Then, for each random sub-sample, we calculated the mean and variance of the predicted edge weights across the samples belonging to each tissue.

**Figure 5.**
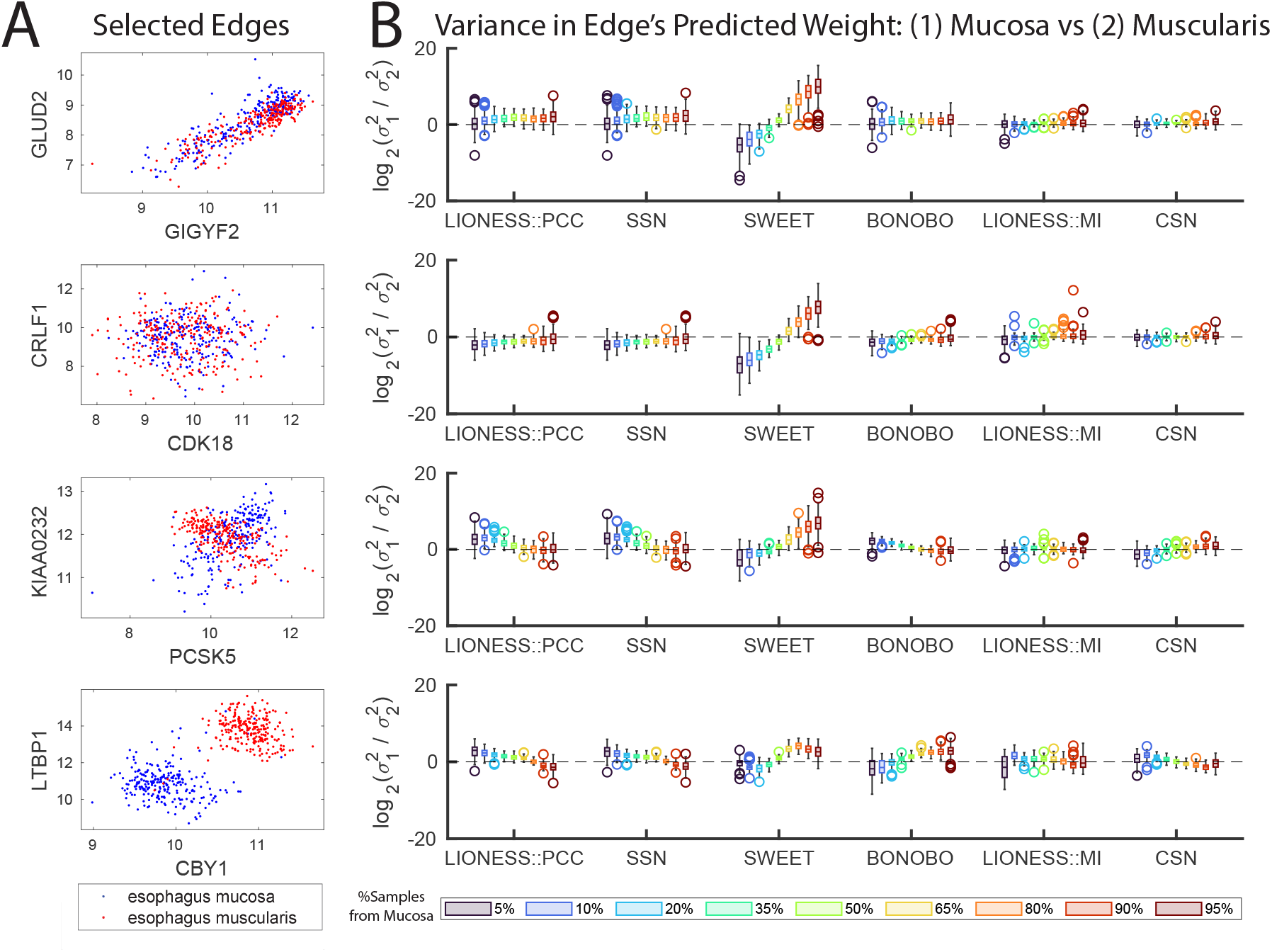
(A) The expression values across all samples in the esophagus muscosa versus esophagus muscularis for four selected edges. (B) For each of the selected edges, the log2-ratio of the variance of the predicted edge weights across samples from the mucosa versus muscularis is shown as a function of the percentage of mucosa samples in the input data (random subset) that was used to infer the edge weights. Box plots show the distribution of this log2-ratio across 100 randomizations. For each box, the bottom, central line, and top represent the 25th percentile, median, and 75th percentiles of the data, respectively; outliers are plotted individually as open circles and whiskers extend to the most extreme data points that are not considered outliers. See also Supplemental Figures 3-4.

The calculated mean edge weights (Supplemental Figure 4) exhibit patterns that are consistent with those observed previously in Figure 2. For the first two exemplar edges (no differential expression or differential correlation), all methods find minimal differences in the mean edge weight across samples from the two tissues, with only subtle trends associated with population size. For the third exemplar edge (no differential expression but high differential correlation), we find that the edge weights predicted by BONOBO and SWEET are highly dependent on the ‘background correlation’, which changes depending on the ratio of samples from the mucosa versus muscularis. However, LIONESS::PCC, SSN, and BONOBO all predict significantly higher edge weights in the mucosa compared to the muscularis, consistent with the known differential correlation. In contrast, SWEET predicts only a small increase in edge weights. Notably, the non-linear methods (LIONESS::MI, CSN) tend to predict higher edge weights for samples from the larger population. This is due to these methods identifying a strong pattern within a subset of samples (those from the larger population) and then down-weighting the remaining samples. Finally, for the fourth exemplar edge (high differential expression but no differential correlation), all methods predict higher edge weights for samples from the smaller population. Like the third exemplar edge, this difference is much smaller for SWEET.

Complementing this analysis, Figure 5 shows the log of the ratio between the variance in edge weights across samples from each tissue. For the first two exemplar edges, the only method that shows large differences in edge weight variability based on population size is SWEET; for these two exemplar edges, the edge weights predicted by SWEET have significantly more variability across samples from the larger population (Figure 5B, top two rows). For the third exemplar edge (Figure 5B, third row), we observe the same trend for SWEET. However, in this case LIONESS::PCC, SSN, and BONOBO also have a subtle trend in the opposite direction, with slightly increased variability in edge weights across samples from the smaller population. Finally, for the fourth exemplar edge (Figure 5B, fourth row), LIONESS::PCC and SSN once again have a slight trend for more variability in predicted edge weights across samples in the smaller population, while SWEET and BONOBO display the opposite trend, with increased variability in predicted edge weights from samples in the larger population. The non-linear methods (LIONESS::MI, CSN) did not display any obvious differences in variability across the predicted edge weights based on subpopulation size.

Our analyses demonstrate that subpopulation structure can impact both the mean and the distribution of predicted sample-specific edge weights. However, the way in which this manifests is different for the different methods and even different edge types, with SWEET, in particular, showing extreme differences in edge weight variability across samples from different subpopulations, including in cases where there is no corresponding differential correlation. This behavior can be traced to the *S*_*q*_ parameter in SWEET, which is consistently lower for samples from the smaller subpopulation, tightening the distribution of predicted edge weights for these samples and resulting in less edge weight variability. This is important for network prediction since higher edge weight variability across samples from one tissue compared to another can lead to a greater percentage of samples from that tissue having a predicted edge weight above a given threshold. This is especially true when the edge weight is similar in both tissues, as is the case for the first, second, and fourth exemplar edges.

## 7 The Challenge of Benchmarking

The analyses above have primarily focused on how various method parameters influence edge weight predictions by characterizing the behavior of individual edges across samples. However, it is also important to evaluate method predictions across all edges in an individual sample, especially since this is what defines a single-sample network.

There is no ideal benchmark for single-sample networks, especially in the context of real-world data [7]. However, we posit that a network for a sample from a given biological context should (1) be similar to another network from that same context and (2) be more similar to another network from that same context than to a network from a different context. In other words, if given a sample from heart tissue, we would expect both that the network inferred for that sample should be similar to an overall heart tissue network (‘accurate’) and *more* similar to an overall heart tissue network than, for example, an overall lung tissue network (‘specific’).

With this in mind, we identified a dataset and developed a corresponding benchmarking approach to evaluate the ‘accuracy’ and ‘specificity’ of single-sample networks in the context of varying expression and population heterogeneity. First, as illustrated in Supplemental Figure 5, we selected 3750 samples from GTEx, 250 each from fifteen different tissues. We then identified three different sets of genes for which to estimate and evaluate sample-specific networks: (1) the 500 genes with the lowest expression variance across these samples, (2) a set of 500 random genes, and (3) the 500 genes with the highest expression variance across these samples. These sets of genes represent data with varying levels of expression heterogeneity. The 500 genes with the lowest expression variance are highly homogeneous, with no discernible differential expression across tissues, while the 500 genes with the highest expression variance have highly distinct expression levels across tissues. The 500 random genes straddle these two extremes, and include genes that exhibit both homogeneous (e.g. housekeeping genes) and heterogeneous (e.g. tissue or subtype-specific genes) expression levels.

Next, we used Pearson correlation to create a reference network for each of the 15 tissues based on these three sets of genes (Figure 6A); we consider these as the underlying network for each of the samples in the tissue. Across these associated tissue-specific reference networks we observe both common structures, with many gene pairs consistently strongly positively or negatively correlated across tissues, as well as tissue-specific patterns. This is true both when there is little to no differential expression of genes across the tissues (low variance) as well as when there is a high level of differential expression of genes across the tissues (high variance), reflecting the fact that differential correlation can occur both with and without significant differential expression.

**Figure 6.**
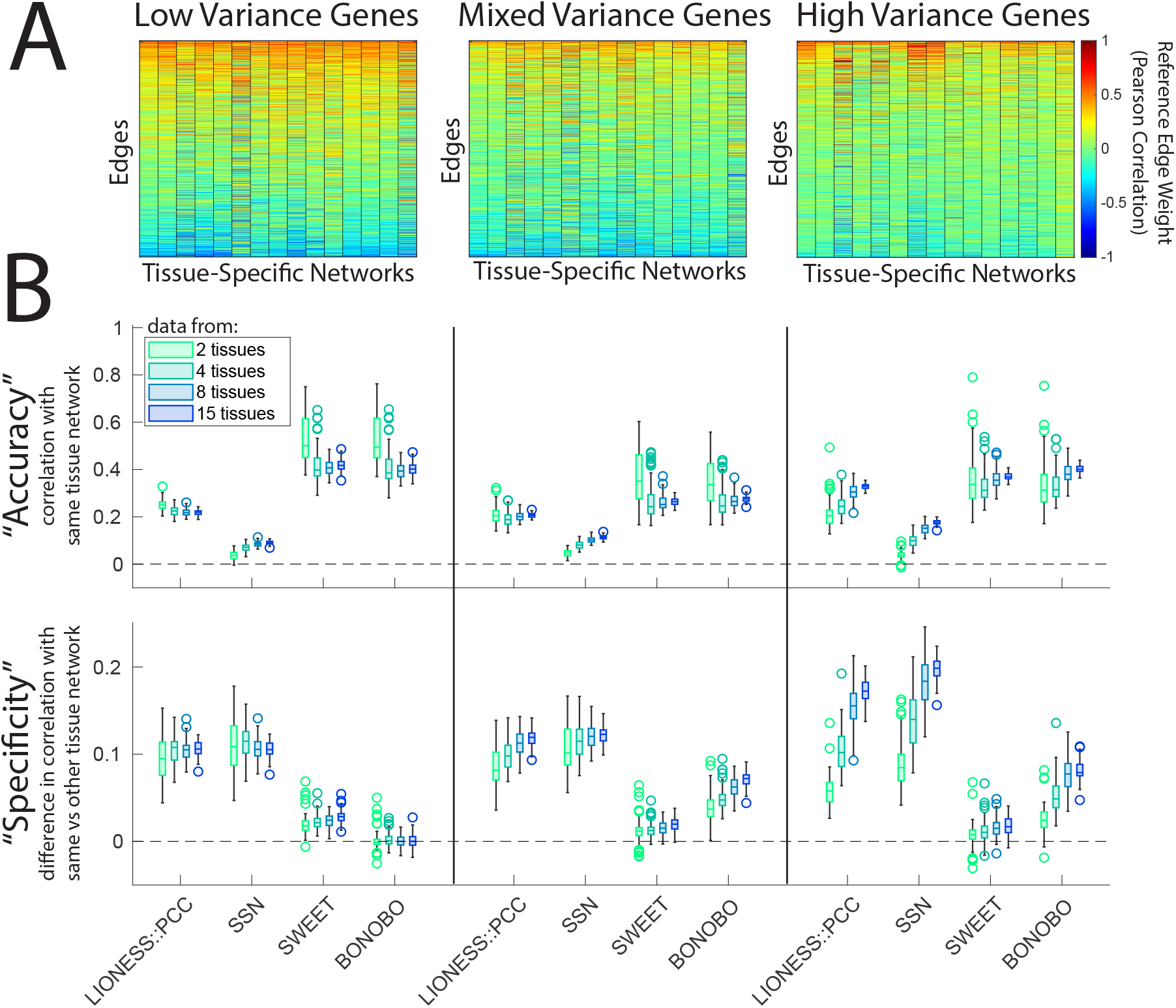
(A) Reference networks (edge weights based on Pearson correlation) for fifteen tissues and between three distinct groups of genes, (1) 500 genes with the lowest variance in expression across samples, (2) 500 randomly selected genes which have mixed variance across samples, and (3) 500 genes with the highest variance in expression across samples. Edges in each plot are ordered based on their mean value across the fifteen tissues. We see both common and tissue-specific edge weight (Pearson correlation) values for all three gene sets. (B) The ‘accuracy’ and ‘specificity’ of the edge weights predicted by the single-sample network methods for each of the three groups of genes when varying the number of tissues (i.e. distinct subpopulations) present in the input data. ‘Accuracy’ was quantified by the mean Pearson correlation of the predicted single-sample networks with their correct associated tissue network (*R*_*t*_; a value of 1 represents perfect correlation and 0 represents random correlation with the correct associated tissue network) ‘Specificity’ was quantified by the mean of the difference between the Pearson correlation of the predicted single-sample networks with their correct associated tissue network and the Pearson correlation of the predicted networks with another tissue network (*R*_*t*_ − *R*_*o*_; a value of 1 represents perfect correlation with the correct tissue network and no correlation with the other tissue network, a value of 0 represents similar correlation with both the correct and other tissue network). To create distributions of values, the analysis was repeated 100 times for different sets of samples. For each box, the bottom, central line, and top represent the 25th percentile, median, and 75th percentiles of the data, respectively; outliers are plotted individually as open circles and whiskers extend to the most extreme data points that are not considered outliers. See also, Supplemental Figure 5-7.

Next, we applied each of the four Pearson-based sample-specific network approaches (LIONESS::PCC, SSN, SWEET, BONOBO) to random sets of samples that represent different levels of population heterogeneity. To create a random set of samples, we randomly selected *T* tissues, determined all 250 × *T* associated gene expression samples, and then randomly selected 100 of these associated samples. Each of these randomly selected samples was then paired both with its correct reference tissue network as well as an ‘other’ tissue network, which was randomly selected from among one of the other *T* − 1 tissues. Next, we applied each single-sample network method to the gene expression data for the 100 selected samples. We then determined the similarity (as measured by Pearson correlation) of each predicted single-sample network with (1) its associated correct tissue network (*R*_*t*_) and (2) its randomly assigned ‘other’ tissue network (*R*_*o*_). We used these two values to determine the ‘accuracy’ (quantified through *R*_*t*_) and ‘specificity’ (quantified through *R*_*t*_ − *R*_*o*_) of each predicted sample-specific network; based on these measures, ideal accuracy/specificity would have a value close to 1 while ‘random’ accuracy/specificity would have a value close to 0. A schematic showing this process is illustrated in Supplemental Figure 6. We repeated this procedure 100 times for *T* equal to 2, 4, 8, and 15 tissues, as well as for the low, random, and high variance gene sets. Varying the number of tissues, *T*, determines the number of known distinct subpopulations in the data while the different gene sets determine the amount of heterogeneity in expression levels across those subpopulations. The results of this analysis are shown in Figure 6B. A similar analysis for the non-linear single sample network methods (LIONESS::MI, CSN) is shown in Supplemental Figure 7.

We observe that the networks predicted by BONOBO and SWEET are generally the most ‘accurate,’ the networks predicted by SSN are the least ‘accurate,’ and the networks predicted by LIONESS::PCC tend to fall in between. On the other hand, the networks predicted by SSN are generally the most ‘specific,’ followed closely by the networks predicted by LIONESS::PCC, the networks predicted by SWEET tend to be the least ‘specific,’ and the networks predicted by BONOBO tend to fall in between. This distinct contrast between the accuracy and specificity of the methods can be understood by recognizing that, as seen in Figure 6A, there are many similarities across the reference tissue-specific networks (in addition to subtle differences). Therefore, it is possible for a network to be highly accurate by capturing only shared network patterns (and not the tissue-specific patterns). Such a highly ‘accurate’ network would have little to no specificity, as it would be no more similar to its correct reference tissue network compared to any other random tissue network. In the context of sample-specific networks, ‘specificity’ is at least, if not more important than ‘accuracy’, since the stated goal of these methods is to infer a network that is specific to a given sample, not a network that represents the larger population of samples. We point out that the results shown here are consistent with the mathematical formulation of these methods (Figure 1). As we have observed in previous sections, both SWEET and BONOBO include a scale factor that pushes their predicted edge weights towards the ‘background correlation/covariance’ term, which captures shared network structures rather than sample-specific (tissue-specific) patterns. At the same time, the SSN equation does not include a ‘background correlation’ term, which improves its ability to capture sample-specific (tissue-specific) structure but minimizes its ability to capture shared network patterns.

Finally, we note that the ‘accuracy’ and ‘specificity’ of the predicted single-sample networks differs based on the variance of genes and the number of subpopulations (tissues) in the data. For networks computed based on low variance genes, there are small shifts in the ‘accuracy’ depending on the number of tissues, with SSN having slightly increased ‘accuracy,’ and LIONESS::PCC, SWEET, and BONOBO having slightly decreased ‘accuracy’ with more tissues. In addition, although networks predicted by SWEET generally have the lowest ‘specificity,’ when using data from low variance genes, the ‘specificity’ of BONOBO is essentially zero, implying that, in this context, these networks contain absolutely no information that is specific to individual samples, but only information that is shared across samples. In contrast, for networks computed based on high variance genes, the ‘accuracy’ of the predicted networks increases moderately for all methods with more tissues. In addition, all methods except for SWEET see a very large increase in ‘specificity’ with more tissues. As expected, the trends for random genes fall between those of the low variance genes and the high variance genes.

Although there are some clear trends, this analysis demonstrates that heterogeneity in both the expression levels (low versus high variance genes) and the number of distinct subpopulations (varying number of tissues) in a dataset can impact the perceived performance of single-sample network methods. This highlights how challenging it can be to benchmark and compare these methods, and ultimately, to determine which method is most appropriate for individual future applications.

## 8 Discussion

In this paper, we explore the mathematical formulation, parameters, and associated outputs of five single-sample network methods. This includes four approaches that have either been explicitly derived in the context of, or can be applied to, Pearson correlation (LIONESS::PCC, SSN, SWEET, BONOBO), and two ‘non-linear’ approaches (LIONESS::MI, CSN). Re-casting the mathematics of the four linear single-sample network approaches into common variables allowed us to identify many synergies as well as key differences. By applying these approaches to both toy and ‘real-world’ gene expression data, we provide an intuition as to how method-specific parameters can impact single-sample network prediction. Our results demonstrate the importance of developing an intuition for these methods grounded in their mathematical formulation.

Although the single-sample edge weights predicted by the four linear methods were generally highly correlated, our analyses showed that the nature of that correlation was often context and parameter-dependent. For example, LIONESS depends on an underlying network reconstruction method, such as Pearson correlation (LIONESS::PCC) or MI (LIONESS::MI), while SSN computes a statistic (Z-score) that characterizes how much a sample’s network differs from that of a reference population. Despite these conceptual differences, the mathematical formulas for LIONESS::PCC and SSN are very similar (see Figure 1) and we find that the edge weights predicted by these two methods are always perfectly correlated, differing only in terms of their magnitude. In contrast, the formulas for SWEET and BONOBO include sample-specific scale factors that depend on all entities (e.g. genes) in the input data. Our analyses demonstrate that these scale factors reduce the impact of the differential correlation/covariance term in the SWEET and BONOBO equations. The result is that the edge weights predicted by these methods are always extremely close to the ‘aggregate’ Pearson correlation network.

In SWEET, the stated goal of the sample-specific scale factor (*K* × *S*_*q*_) is to account for potential biases due to differences in subpopulation size [29]. We find that the value of this scale factor is consistently lower for samples from smaller subpopulations, which tightens the distribution of predicted edge weights for these samples, leading to consistently less edge weight variability. The corresponding increased variability in edge weight across samples from the larger of two subpopulations can lead to a bias for an increased proportion of those samples to have edge weights above a given threshold. In contrast, we find that the value of the sample-specific scale factor used by BONOBO 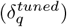 does not vary significantly across samples, but instead depends on the heterogeneity of the input data. In addition, this scale factor can be unintentionally impacted by how the input data are preprocessed and/or normalized. For example, data preprocessing that results in either a very high average variance or a highly consistent variance across all genes can lead to 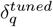 approaching zero, driving all edge weights to the ‘aggregate’ Pearson correlation. The consequence of this can be observed in our toy data analysis, where we found that using the default of 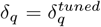 resulted in BONOBO’s predictions being *negatively* correlated with the predictions of all other tested methods. With this in mind, it might be beneficial for a user to manually set *δ*_*q*_ = *δ*, a constant.

The single-sample edge weights predicted by the two non-linear methods (LIONESS::MI, CSN) were also generally correlated; however, the edge weights predicted by the non-linear compared to the linear methods were often distinct. For example, in the toy data analysis, we observed that both non-linear methods gave high weights to all of the samples that made up the X pattern of the non-linear edge, while the linear methods generally gave higher weights to samples that were along the *x* = *y* diagonal. Given these caveats, we suggest that future studies that use these methods evaluate the underlying data from which any interesting edges are derived, as we did in Figure 2. One strength of LIONESS is that it can be applied to different underlying aggregate network reconstruction approaches [20]. In fact, it is commonly used to model sample-specific gene regulatory networks (rather than correlation networks) [27, 28, 25]. However, naively applying LIONESS can have drawbacks. For example, although we found similar results using LIONESS::MI and CSN, CSN was much more computationally efficient and consistently ran much faster than LIONESS::MI. This is because CSN is specifically optimized for identifying a sample-specific non-linear relationship, whereas LIONESS is an equation that can be applied to MI, but does not directly incorporate any information about how MI is calculated. LIONESS has previously been optimized for application to Pearson correlation [35, 19] and ARACNe [25]; optimization for application to MI may be a promising future direction.

Our analysis also simultaneously underlines why it can be dangerous to rely too much on simulated data *and* why it can be powerful to use simulated data for method evaluation. In particular, the very low values we observed for 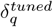 and the population substructure we uncovered when evaluating *S*_*q*_ were not in the toy data by design. Rather, these were unintended features that were uncovered through analysis and subsequently traced back to how we generated these data. However, because these unintended features were distinct and easily quantifiable, we were able to use these findings to better understand how *S*_*q*_ and 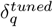 interact with specific data features. Had we only relied on real data, complex and overlapping substructures within the data may have prevented us from clearly characterizing these parameters. In Figure 5 we show that differential expression, differential correlation, and differences in subpopulation size all can influence predicted edge weights. Other features not explored here, such as differential variability, overlapping subpopulations, and/or data sparsity, may also impact single-sample network predictions. Even the ‘real-world’ expression data we used in our analyses has assumed structural features that may unintentionally impact edge weight predictions. For example, these data contained an equal number of samples from each tissue and only genes that are expressed across all tissues (see Methods). Nevertheless, we believe that these data are consistent with the underlying assumptions of BONOBO and SWEET, which was necessary for a fair assessment of *S*_*q*_ and 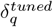.

Our final analysis explored the impact of data heterogeneity on single-sample network prediction. Across different levels of both expression and subpopulation heterogeneity, our results identified a critical trade-off when trying to simultaneously predict network edges that are shared across samples as well as those that are specific to a given sample. In particular, we found that BONOBO and SWEET, whose predicted networks are highly similar to the ‘aggregate’ network, consistently estimated the most accurate but least specific networks, while SSN estimated the most specific but least accurate networks. LIONESS::PCC straddled these domains, with an accuracy almost as high as SWEET and BONOBO and a specificity almost as high as SSN in most evaluations. The output of a single-sample network algorithm is typically either directly analyzed to identify biological signals or used as an input to another downstream algorithm. The trade-off between predicted network accuracy and specificity is critical to consider in this larger context. A primary goal of sample-specific network modeling is to obtain a network that is *specific* to a given sample. If specificity is not needed or desired, our results suggest that it may be more effective to analyze an ‘aggregate’ network, rather than to use SWEET or BONOBO to generate a set of non-specific sample-specific networks.

These results also highlight how challenging it can be to benchmark single-sample network methods. We find that how one designs the benchmark data and selects the genes, edges, and performance metric, all can impact perceived method performance. Recognizing these issues is especially relevant for those who work in method development, where showing superior performance, at least in a particular context, is often necessary for publication. In addition, describing the potential caveats and limitations of a new method can sometimes lead to unwelcome reviewer comments. This ecosystem encourages papers that emphasize and exploit the perceived limitations of existing methods and leads to comparative analyses that place the new method in the most positive light. This can lead to biases in the published literature.

An additional complicating factor is that it can sometimes be challenging to run other methods, as they tend to be programmed in different languages with different levels of documentation. It also takes significant effort to understand various methods’ assumptions, parameters, and inner-workings, and to potentially re-implement them in a consistent fashion. However, our study illustrates that failing to do this can lead to inadvertently missing important synergies between existing methods. Along these lines, we believe that our work points to important opportunities for the field of single-sample network inference. We advocate for new methods, both in this area and in other areas of computational biology, to strive to compare with existing approaches using a unified mathematical terminology, such as what we show in Figure 1. Reviewers should also commend efforts by authors to demonstrate both the synergies and differences between new and existing approaches, and encourage the publication of results that show both where a new method succeeds and where it does not. Although doing this will take significant effort by both authors and reviewers, we believe that, together, the community will be stronger if this can be accomplished.

## 9 Methods

### 9.1 Single-sample network methods

#### 9.1.1 Overview of Single-sample Network Methods

In this paper, we explore six single-sample network methods in detail. This includes four ‘linear’ methods which were either explicitly derived in the context of Pearson correlation or can be applied to Pearson correlation - LIONESS::PCC, SSN, SWEET, BONOBO - and two ‘non-linear’ methods - LIONESS::MI, CSN. A detailed summary of the mathematics underlying each of these six methods is provided in Supplemental File 1.

#### 9.1.2 Implementing the Single-sample Network Methods

For our primary analysis, we implemented the single-sample network methods based on the mathematical framework presented in Figure 1 and described in detail in Supplemental File 1. Since this recasting sometimes involved several steps, as we implemented each method, we verified that our mathematical recasting was correct by confirming that the new implementation of the method either exactly reproduced the original implementation provided by the authors of the method [36, 37, 38, 39] and/or a key result reported for that method in a publication that used the original implementation. Parameters in the main implementation were kept simple, and matched the default parameters in the original method papers/implementations, including *K* = 0.1 and *ϵ* = 0.01 for SWEET, 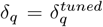 for BONOBO, and boxsize=0.1 for CSN. For LIONESS::MI we leveraged an MI implementation available at https://github.com/otoolej/mutual_info_kNN. We note that the original CSN implementation by default filters out any edges with weight less than zero. In contrast, our implementation reports values for all edges and all samples (which is an option in the original CSN implementation). The original CSN implementation also assumes non-negative input data. Therefore, we ensured that the data used in our analyses did not contain negative values. However, within our implementation of CSN we include a comment that explains how the implementation could be adjusted to handle negative data.

We also created several additional implementations of these methods that allowed us to systematically evaluate parameters (such as *K* or *δ*_*q*_ for SWEET and BONOBO, respectively), optimize the computation in a particular setting (e.g. calculating the weights for a single edge rather than all edges), or to evaluate the impact of method conceptualization (e.g. to frame the methods as ‘add-one-in’ instead of ‘leave-one-out’, see Supplemental File 3). All implementations are provided in our github repository: https://github.com/kimberlyglass/single-sample-networks/tree/main/NetworkCode.

### 9.2 Expression Data

#### 9.2.1 Toy Data

To create the toy expression data, we used a normal random distribution function to create a vector (*v*) of length *L* = 250, with a mean *µ* = 0, and a standard deviation *σ* = 1, and sorted in increasing order. We replicated this vector multiple times to create a matrix (*M*) with six rows and 500 columns that contained values that defined two populations (sample set 1 and sample set 2) and two distinct edge types (*M* = [*v, v, v, v, v, v*; *v, v, v*, −*v*, −*v*, −*v*]). We then used a normal random distribution function (*µ* = 0, *σ* = 0.1) to add a small amount of noise to this matrix. We created an additional population (sample set 3 / noise) by using a normal random distribution function (*µ* = 0, *σ* = 1) to create a matrix (*R*) with six rows and 100 columns. We concatenated these two matrices (*D* = [*M, R*]) to create a final matrix with six rows and 600 columns. We then shifted all values in this matrix to be greater than zero *D*_*f*_ = *D* + min[*D*] + *ϵ*^*′*^, where *ϵ*^*′*^ = 2.22 × 10^*−*16^. This toy data is visualized in Supplemental Figure 1A and is available in our github repository: https://github.com/kimberlyglass/single-sample-networks.

#### 9.2.2 GTEx Data

We leveraged gene expression data from GTEx [34, 40] that had been previously processed using YARN [41] and which is described in detail in [5]. From these data, we selected fifteen tissues that contained at least 250 associated samples. This included samples from the subcutaneous adipose, tibial artery, three different brain regions, fibroblasts, esophagus mucosa, esophagus muscularis, heart left ventricle, lung, skeletal muscle, tibial nerve, skin, thyroid, and whole blood. For each tissue, we selected 250 random samples and calculated the median expression of genes across those samples. We then filtered to only include protein-coding genes that had a median expression of at least 2 in all tissues, resulting in a final data-set with 15 × 250 = 3750 samples and 13281 genes. For the analyses shown in Figures 4-5 we leveraged the 500 samples from these data that were associated with either the esophagus muscularis or esophagus mucosa. For the analyses shown in Figure 6 we leveraged all 3750 samples from these data. These data are visualized in Supplemental Figure 5.

### 9.3 Data and Code availability

All data and code necessary to reproduce the analyses and figures in this manuscript are available at https://doi.org/10.5281/zenodo.18509155 and https://github.com/kimberlyglass/single-sample-networks. A python package that implements the four ‘linear’ single-sample network methods based on their re-cast formulation is available at https://github.com/kimberlyglass/single-sample-networks/tree/main/python.

## Supporting information

Supplemental File 1 -- Supplemental Derivations

Supplemental File 3 -- Supplemental Analyses

## 9.4 Acknowledgments

We wish to thank Jeremie Fish for providing feedback on the MI implementation used for LIONESS::MI and John Quackenbush and Federico Melograna for providing feedback on the manuscript draft.

