## Supplemental File 1 -- Supplemental Derivations for "Challenges and Opportunities in Single-Sample Network Modeling"

### Supplemental File 1:
### Single-Sample Network Method (re)Derivations

Marieke Kuijjer, Margherita De Marzio, & Kimberly Glass

February 27, 2026

#### 1 Generalizable methods for estimating single-sample networks

##### 1.1 LIONESS

LIONESS (Linear Interpolation to Obtain Network Estimates for Single Samples) is a method-agnostic mathematical formula to estimate single-sample networks. This formula is defined in Equation 4 of Kuijjer et al. [1]:

$$e_{ij}^{(q)} = \frac{1}{w_q}(e_{ij}^{(\alpha)} - e_{ij}^{(\alpha-q)}) + e_{ij}^{(\alpha-q)} \tag{1}$$

where $e_{ij}^{(q)}$ is the weight of an edge from $i$ to $j$ in the network for a specific sample $q$ in a dataset, $e_{ij}^{(\alpha)}$ is the weight of that same edge in a network reconstructed using all samples in the dataset, and $e_{ij}^{(\alpha-q)}$ is the weight of that same edge in a network reconstructed using all samples in the dataset except sample $q$. $w_q$ is the weight given to the individual sample $q$ when reconstructing the $e_{ij}^{(\alpha)}$ network, with the stipulation that the sum of $w_q$ across all samples equals 1, i.e. $\Sigma_{k=1}^{N} w_k = 1$, where $N$ is the total number of samples in the dataset. This means that in cases where all samples are weighted equally, $w_q = \frac{1}{N}$. The networks $e_{ij}^{(\alpha)}$ and $e_{ij}^{(\alpha-q)}$ are calculated using an external 'aggregate' network reconstruction approach, with Pearson correlation, Mutual Information, PANDA, and CLR shown as exemplar approaches in Kuijjer et al. [1].

#### 2 Estimating single-sample networks based on Pearson correlation

##### 2.1 LIONESS::PCC

In this paper, we recast LIONESS for application to networks constructed using Pearson correlation coefficient (referred to as LIONESS::PCC). In this case, the following variable substitutions can be made: (1) $e_{ij}^{(q)} \sim \rho_{ij}^{(q)}$, the estimated sample-specific network, (2) $e_{ij}^{(\alpha)} \sim \rho_{ij}^{(\alpha)}$, the Pearson correlation between two entities (e.g. genes), $i$ and $j$, across all samples, and (3) $e_{ij}^{(\alpha-q)} \sim \rho_{ij}^{(\alpha-q)}$, the Pearson correlation between two entities, $i$ and $j$, across all samples except for $q$. Pearson correlation gives each sample equal weight (although a weighted form exists, see Section 3.1 below), so $w_q = \frac{1}{N}$. Thus, in the context of Pearson correlation, Equation 1 can be recast as:

$$\rho_{ij}^{(q)} = N(\rho_{ij}^{(\alpha)} - \rho_{ij}^{(\alpha-q)}) + \rho_{ij}^{(\alpha-q)}. \tag{2}$$

##### 2.2 SSN

The mathematical formulation of SSN (sample-specific network) is defined in Equation 2 and shown in Figure 2 of Liu et al. [2]:

$$Z = \frac{\Delta PCC_n}{\frac{1-PCC_n{}^2}{n-1}}, \tag{3}$$

where $Z$ is the Z-value (statistic) for edges in a sample-specific network, $\Delta PCC_n = PCC_{n+1} - PCC_n$ is the change in Pearson correlation between a 'reference network' ($PCC_n$) and 'perturbed network' ($PCC_{n+1}$), and $n$ is the number of samples used to calculate the reference network; the reference network, $PCC_n$, is the Pearson correlation coefficient calculated using $n$ samples, and the perturbed network, $PCC_{n+1}$, is the Pearson correlation coefficient calculated using $n+1$ samples, i.e. after adding in an additional sample.

This means that to recast SSN using the same variable names as used for LIONESS::PCC, i.e. for a single-sample $q$ and a single edge (between $i$ and $j$) the following substitutions can be made: $Z \sim \rho_{ij}^{(q)}$, $PCC_{n+1} \sim \rho_{ij}^{(\alpha)}$, $PCC_n \sim \rho_{ij}^{(\alpha-q)}$, and

$n \sim N-1$. Thus, Equation 3 can be recast as follows:

$$\rho_{ij}^{(q)} = \frac{\rho_{ij}^{(\alpha)} - \rho_{ij}^{(\alpha-q)}}{\frac{1-(\rho_{ij}^{(\alpha-q)})^2}{N-2}} \tag{4}$$

This can be equivalently written, with minor rearrangements as:

$$\rho_{ij}^{(q)} = \frac{1}{1-(\rho_{ij}^{(\alpha-q)})^2}(N-2)(\rho_{ij}^{(\alpha)} - \rho_{ij}^{(\alpha-q)}) \tag{5}$$

##### 2.3 SWEET

The SWEET (sample-specific weighted correlation network) method is formulated in Equation 2 of Chen et al. [3]:

$$E_{ij}^{(S_p)} = W^{(S_p)} \times n \times K(E_{ij}^{(n+S_p)} - E_{ij}^{(n)}) + E_{ij}^{(n)} \tag{6}$$

where $E_{ij}^{(n)}$ is the Pearson correlation coefficient calculated for an edge between genes $i$ and $j$ across $n$ samples, referred to in Chen et al. as the 'aggregate network', and $E_{ij}^{(n+S_p)}$ is the Pearson correlation coefficient calculated for an edge between genes $i$ and $j$ across $n+1$ samples, referred to in Chen et al. as the 'perturbed network'. Importantly, the sample of interest ($S_p$) is included twice across the $n+1$ samples used to calculate $E_{ij}^{(n+S_p)}$ – once as a member of the original $n$ samples and once as the additional sample of interest. We note that it is considered inappropriate statistical practice to include the same sample multiple times when calculating the Pearson correlation. Therefore, when re-casting SWEET we opt to instead characterize the 'aggregate' network as one computed across $n-1$ samples (all but the sample of interest) and the 'perturbed' network as the one computed across all $n$ samples (all samples, which includes the sample of interest). We note that this shift impacts the interpretation of $n$ in Equation 6, from representing all samples in the dataset, to representing all but one. However, in the limit of many samples $(n-1) \approx n$. We also note that we have tested implementations of SWEET both with and without this shift in interpretation. The result of this analysis can be found at `https://github.com/kimberlyglass/single-sample-networks/tree/main/Supplemental`. We observe that this shift has no quantifiable impact on the final computed value of $E_{ij}^{(S_p)}$ and absolutely no impact on the final conclusions of our manuscript.

SWEET also defines two additional parameters, a 'balance parameter' $K$ (which is set equal to a constant value, default value is $K = 0.1$), and $W^{(S_p)}$, a 'genome-wide sample weight' (i.e. a weight associated with sample $S_p$). In Chen et al. this sample specific weight, $W^{(S_p)}$, is defined in Equation 1 as:

$$W^{(S_p)} = \frac{\mu_{PCC}^{(S_p)} - \min(\mu_{PCC}^{(S)}) + x}{\max(\mu_{PCC}^{(S)}) - \min(\mu_{PCC}^{(S)}) + x} \tag{7}$$

where $\mu_{PCC}^{(S)}$ is a vector representing the average Pearson correlation of each sample with all other samples and $\mu_{PCC}^{(S_p)}$ is the element of that vector for sample $S_p$; $x$ is a small value, which in the Chen et al. paper is set equal to 0.01. In other words, whereas other references to the Pearson correlation coefficient have quantified the relationship between genes, the sample-specific weight $W^{(S_p)}$ conceptualized in Chen et al. leverages the Pearson correlation coefficient between pairs of *samples*.

To recast $W^{(S_p)}$ using variable names more consistent with other methods, we defined $r_{kq}$ as the Pearson correlation between samples $k$ and $q$ across all genes. Based on this, $\mu_{PCC}^{(S_p)}$ can be recast as $r_q = \frac{1}{N-1}\Sigma_{k\neq q}^{N} r_{kq}$, the average Pearson correlation of sample $q$ with all other samples (except for itself), and $\mu_{PCC}^{(S)}$ can be recast as $r_{.}$, a vector containing these values for all $N$ samples in the dataset. We also recast $x$ as $\epsilon$. Based on this, Equation 7 can be re-written as follows:

$$S_q = \frac{r_q - \min(r_{.}) + \epsilon}{\max(r_{.}) - \min(r_{.}) + \epsilon} \tag{8}$$

Based on this, we see that $S_q$ can take a value between 0 and 1, $S_q \to 0$ for the sample with the lowest average correlation with other samples in the dataset ($r_q = \min(r_{.})$), and $S_q = 1$ for the sample with the highest average correlation with other samples in the dataset ($r_q = \max(r_{.})$).

Finally, to recast SWEET using the same variable names as used for LIONESS::PCC and SSN, i.e. for a single-sample $q$ and a single edge (between genes $i$ and $j$) the following substitutions can be made: $W^{(S_p)} \sim S_q$ (see Equation 7 versus Equation 8), $n \sim N-1$, $E_{ij}^{(n+S_p)} \sim \rho_{ij}^{(\alpha)}$, and $E_{ij}^{(n)} \sim \rho_{ij}^{(\alpha-q)}$. Thus, Equation 6 can be re-written as follows:

$$\rho_{ij}^{(q)} = (K \times S_q)(N-1)(\rho_{ij}^{(\alpha)} - \rho_{ij}^{(\alpha-q)}) + \rho_{ij}^{(\alpha-q)} \tag{9}$$

#### 2.4 BONOBO

The BONOBO (Baysian optimized networks obtained by assimilating omics) approach, as shown in Equation 4 of the Saha et al. [4], estimates the sample-specific covariance matrix as:

$$\Sigma_i = \delta_i(x_i - \bar{x})(x_i - \bar{x})^T + (1 - \delta_i)S_i \tag{10}$$

where $x_i$ is a vector of length $g$ containing the expression of genes in sample $i$, $\bar{x}$ is a vector of length $g$ containing the mean expression of genes across all samples, $S_i$ is a $g \times g$ matrix containing the covariance values between all pairs of genes, calculated across all samples except for sample $i$, and $\delta_i = \frac{1}{\nu_i - g}$ is a hyperparameter with $\nu_i$ equal to the degrees of freedom; $g$ is the number of genes. In the last paragraph of the Methods section in their paper, Saha et al. define the sample-specific correlation between two genes, $j$ and $k$ as:

$$\rho_{jk}^{(i)} = \frac{v_{jk}^{(i)}}{\sqrt{v_{jj}^{(i)} v_{kk}^{(i)}}} \tag{11}$$

where $v_{jk}^{(i)}$ denotes the $(j, k)$ entry in the sample-specific covariance matrix $(V_i)$, which is assumed to be equal to $\Sigma_i$ (equation 10); in other words, $v_{jk}^{(i)}$ denotes the $(j, k)$ entry of $\Sigma_i$.

To recast BONOBO using variable names consistent with those used for LIONESS::PCC, SSN, and SWEET, i.e. for a single-sample $q$, the following substitutions can be made: $x_i \sim x^{(q)}$ (a vector containing the expression value for each gene in sample $q$), $\bar{x} \sim \bar{x}^{(\alpha)}$ (a vector containing the average expression value of each gene across all samples), $S_i \sim \sigma^{(\alpha-q)}$ (the pairwise covariance between all genes across all samples except for $q$), and $\delta_i \sim \delta_q$. Based on this, for a single edge between genes $i$ and $j$, Equation 10 can be re-written as:

$$\sigma_{ij}^{(q)} = \delta_q(x_i^{(q)} - \bar{x}_i^{(\alpha)})(x_j^{(q)} - \bar{x}_j^{(\alpha)}) + (1 - \delta_q)\sigma_{ij}^{(\alpha-q)} \tag{12}$$

or equivalently, with minimal rearrangement:

$$\sigma_{ij}^{(q)} = \delta_q[(x_i^{(q)} - \bar{x}_i^{(\alpha)})(x_j^{(q)} - \bar{x}_j^{(\alpha)}) - \sigma_{ij}^{(\alpha-q)}] + \sigma_{ij}^{(\alpha-q)} \tag{13}$$

We note that in the limit of large $N$, Equation 13 can be further simplified. First, we note that the covariance between two genes, $i$ and $j$ across all samples is defined as:

$$\begin{aligned}
\sigma_{ij}^{(\alpha)} &= \frac{1}{N-1}\sum_{k=1}^{N}(x_i^{(k)} - \bar{x}_i^{(\alpha)})(x_j^{(k)} - \bar{x}_j^{(\alpha)}) \\
&= \frac{1}{N-1}\left[(x_i^{(q)} - \bar{x}_i^{(\alpha)})(x_j^{(q)} - \bar{x}_j^{(\alpha)}) + \sum_{k \neq q}^{N}(x_i^{(k)} - \bar{x}_i^{(\alpha)})(x_j^{(k)} - \bar{x}_j^{(\alpha)})\right] \\
&= \frac{(x_i^{(q)} - \bar{x}_i^{(\alpha)})(x_j^{(q)} - \bar{x}_j^{(\alpha)})}{N-1} + \sigma_{ij}^{(\alpha-q)}
\end{aligned} \tag{14}$$

Rearranging we find that:

$$(x_i^{(q)} - \bar{x}_i^{(\alpha)})(x_j^{(q)} - \bar{x}_j^{(\alpha)}) = (N-1)(\sigma_{ij}^{(\alpha)} - \sigma_{ij}^{(\alpha-q)}) \tag{15}$$

which we can substitute into Equation 13:

$$\begin{aligned}
\sigma_{ij}^{(q)} &= \delta_q[(N-1)(\sigma_{ij}^{(\alpha)} - \sigma_{ij}^{(\alpha-q)}) - \sigma_{ij}^{(\alpha-q)}] + \sigma_{ij}^{(\alpha-q)} \\
&= \delta_q[(N-1)\sigma_{ij}^{(\alpha)} - N\sigma_{ij}^{(\alpha-q)}] + \sigma_{ij}^{(\alpha-q)}
\end{aligned} \tag{16}$$

In the limit of large $N$ this reduces to:

$$\sigma_{ij}^{(q)} = \delta_q N(\sigma_{ij}^{(\alpha)} - \sigma_{ij}^{(\alpha-q)}) + \sigma_{ij}^{(\alpha-q)} \tag{17}$$

We have tested implementations of BONOBO both with the mathematics in Equation 13 and in the limit of large $N$ (Equation 17). The result of this analysis can be found at `https://github.com/kimberlyglass/single-sample-networks/tree/main/Supplemental`. We observe that re-casting the BONOBO equation assuming large $N$ has absolutely no impact on the final conclusions of our manuscript. Thus for our main implementation, we use the form of BONOBO shown in Equation 17.

Based on this recast definition of the sample-specific covariance, $\sigma_{ij}^{(q)}$ (see Equations 13 and 17), we can recast BONOBO's definition of the sample-specific Pearson correlation (see Equation 11) as:

$$\rho_{ij}^{(q)} = \frac{\sigma_{ij}^{(q)}}{\sqrt{\sigma_{ii}^{(q)}\sigma_{jj}^{(q)}}} \tag{18}$$

The BONOBO equation also has a sample-specific tuning parameter, $\delta_i$ (see Equation 10) which we recast above to $\delta_q$. According to Saha et al., this tuning parameter can be manually set equal to a constant value between 0 and 1, i.e. $\delta_q = \delta$ for all samples. However, by default this tuning parameter is set equal to a sample-specific value, which is defined in Equation 8 of Saha et al. as:

$$\delta_i = \frac{1}{\nu_i - g} = 1/\left[3 + \frac{2\sum_{k=1}^{g}(s_i^{(kk)})^2}{\sum_{k=1}^{g}\eta^{(k)}}\right]. \tag{19}$$

When recasting the equation for $\delta_i$, we both examined the mathematics in Saha et al. and compared directly with the implementation of BONOBO on netZoo [5] (access date: January 16, 2026; see lines 60-64) to ensure the validity of our interpretation. Based primarily on the provided implementation of BONOBO, we make the following substitutions to recast $\delta_i$ as $\delta_q^{tuned}$: (1) $\sum_{k=1}^{g}(s_i^{(kk)})^2 \sim \text{mean}[\sqrt{\sigma_{\cdot}^{(\alpha-q)}}]$ and (2) $\sum_{k=1}^{g}\eta^{(k)} \sim \text{var}[\sigma_{\cdot}^{(\alpha-q)}]$, where $\sigma_{\cdot}^{(\alpha-q)}$ is a vector containing all $\sigma_{ii}^{(\alpha-q)}$, i.e. the diagonal entries of $\sigma^{(\alpha-q)}$ for all genes. Thus, Equation 19 can be re-written as follows:

$$\delta_q^{tuned} = 1/\left[3 + \frac{2 \times \text{mean}[\sqrt{\sigma_{\cdot}^{(\alpha-q)}}]}{\text{var}[\sigma_{\cdot}^{(\alpha-q)}]}\right]. \tag{20}$$

Examination of this equation shows that $\delta_q^{tuned}$ can vary between zero (when the mean of $\sigma_{\cdot}^{(\alpha-q)}$ is much larger than the variance of $\sigma_{\cdot}^{(\alpha-q)}$) and 1/3 (when the mean of $\sigma_{\cdot}^{(\alpha-q)}$ is much smaller than the variance of $\sigma_{\cdot}^{(\alpha-q)}$).

#### 2.5 Summary

Through recasting the variables from each of the methods described above, we can identify many mathematical similarities and better delineate mathematical differences. To better emphasize these similarities and differences, we summarize the methods here, repeating the final forms of the recast equations shown above.

An initial set of variables can be defined, which includes: (1) $\rho_{ij}^{(\alpha)}$, the Pearson correlation between two genes (or other entities), $i$ and $j$, across all samples ($\alpha$); (2) $\rho_{ij}^{(\alpha-q)}$, the Pearson correlation between two genes (or other entities), $i$ and $j$, across all samples except for sample $q$ ($\alpha - q$); and (3) $N$, the total number of samples in the dataset.

Based on these variables, when applied to Pearson correlation, LIONESS::PCC can be written as follows:

$$\rho_{ij}^{(q)} = N(\rho_{ij}^{(\alpha)} - \rho_{ij}^{(\alpha-q)}) + \rho_{ij}^{(\alpha-q)}. \tag{21}$$

SSN, which was explicitly derived with Pearson correlation in mind, can be written as follows:

$$\rho_{ij}^{(q)} = \frac{1}{1-(\rho_{ij}^{(\alpha-q)})^2}(N-2)(\rho_{ij}^{(\alpha)} - \rho_{ij}^{(\alpha-q)}) \tag{22}$$

SWEET requires defining several additional variables, including (1) $K$, a constant that represents a percentage and can vary from 0 to 1. $K$ is set equal to 0.1 in the Chen et al. paper; and (2) $S_q$, a variable that represents sample $q$'s weight and varies between 0 and 1:

$$S_q = \frac{r_q - \min(r_{\cdot}) + \epsilon}{\max(r_{\cdot}) - \min(r_{\cdot}) + \epsilon} \tag{23}$$

where $r_q = \frac{1}{N-1}\Sigma_{k\neq q}^{N} r_{kq}$ represents the average Pearson correlation of sample $q$ with all other samples (except itself); $r_{kq}$ denotes the Pearson correlation between samples $k$ and $q$ across all genes (or entities); $r_{\cdot}$ represents a vector containing the$r_q$ values for all $N$ samples in the dataset; $\epsilon$ is a small constant value, which is set equal to 0.01 in the Chen et al. paper.

After defining $K$ and $S_q$, SWEET can be written as follows:

$$\rho_{ij}^{(q)} = (K \times S_q)(N-1)(\rho_{ij}^{(\alpha)} - \rho_{ij}^{(\alpha-q)}) + \rho_{ij}^{(\alpha-q)} \tag{24}$$

BONOBO uses a slightly different, but related, set of input variables, which includes: (1) $\sigma_{ij}^{(\alpha)}$, the covariance between two genes (or other entities), $i$ and $j$, across all samples ($\alpha$); and (2) $\sigma_{ij}^{(\alpha-q)}$, the covariance between two genes (or other entities), $i$ and $j$, across all samples except for sample $q$ ($\alpha - q$). It also defines an additional variable, $\delta_q$, which by default is set equal to:

$$\delta_q^{tuned} = 1/\left[3 + \frac{2 \times \text{mean}[\sqrt{\sigma_{\cdot}^{(\alpha-q)}}]}{\text{var}[\sigma_{\cdot}^{(\alpha-q)}]}\right], \tag{25}$$

a sample-specific 'tuned' value that can vary between 0 and 1/3. Saha et al. state that $\delta_q$ can alternatively be manually set equal to a constant value between 0 and 1 for all samples, i.e. $\delta_q = \delta$. Based on the above, BONOBO defines a sample-specific correlation network by first estimating the sample-specific covariance as:

$$\sigma_{ij}^{(q)} = \delta_q[(x_i^{(q)} - \bar{x}_i^{(\alpha)})(x_j^{(q)} - \bar{x}_j^{(\alpha)}) - \sigma_{ij}^{(\alpha-q)}] + \sigma_{ij}^{(\alpha-q)} \tag{26}$$

where $x_i^{(q)}$ is the value of entity (or gene) $i$ in sample $q$, and $\bar{x}_i^{(\alpha)}$ is the average value of $i$ across all samples ($\alpha$). In the limit of large $N$, Equation 26 can be approximated as:

$$\sigma_{ij}^{(q)} = \delta_q N(\sigma_{ij}^{(\alpha)} - \sigma_{ij}^{(\alpha-q)}) + \sigma_{ij}^{(\alpha-q)} \tag{27}$$

A sample-specific correlation network is then calculated as:

$$\rho_{ij}^{(q)} = \frac{\sigma_{ij}^{(q)}}{\sqrt{\sigma_{ii}^{(q)}\sigma_{jj}^{(q)}}} \tag{28}$$

### 3 Estimating single-sample networks based on weighted Pearson correlation

#### 3.1 LIONESS::WPCC

In the main text of our paper, we apply LIONESS to networks constructed using the standard Pearson correlation coefficient (LIONESS::PCC, see Section 2) as well as MI (LIONESS::MI, see below). However, LIONESS can also be applied to networks calculated using a *weighted* Pearson correlation (referred to as LIONESS::WPCC). In this case, the following variable substitutions can be made: (1) $e_{ij}^{(q)} \sim \varrho_{ij}^{(q)}$, the estimated sample-specific network, (2) $e_{ij}^{(\alpha)} \sim \varrho_{ij}^{(\alpha)}$, the weighted Pearson correlation between two entities (e.g. genes), $i$ and $j$, across all samples, and (3) $e_{ij}^{(\alpha-q)} \sim \varrho_{ij}^{(\alpha-q)}$, the weighted Pearson correlation between two entities, $i$ and $j$, across all samples except for $q$. Thus, in the context of Pearson correlation, Equation 1 can be recast as:

$$\varrho_{ij}^{(q)} = \frac{1}{w_q}(\varrho_{ij}^{(\alpha)} - \varrho_{ij}^{(\alpha-q)}) + \varrho_{ij}^{(\alpha-q)}. \tag{29}$$

Note that in this equation, each sample is given a unique weight ($w_q$), which corresponds to the weight given to that sample when calculating the weighted Pearson for all samples ($\varrho_{ij}^{(\alpha)}$). Although not needed to calculate $\varrho_{ij}^{(q)}$, we note that this equation assumes that samples are given a proportionally equivalent weight when calculating $\varrho_{ij}^{(\alpha-q)}$, i.e. $w_q = Cw_q^{(\alpha-q)}$, where $C$ is a constant. Since $\sum_k^N w_k = \sum_{k\neq q}^N w_k^{(\alpha-q)} = 1$, $C = 1 - w_q$, and $w_q^{(\alpha-q)} = w_q/(1-w_q)$.

Given a dataset with $P$ distinct and known populations, each with a known size $N_p$, one reasonable approach would be to give each *population* equal weight (rather than each individual sample). In this case, a sample $q$ that is within a population $p$ would have a weight $w_q = \frac{1}{PN_p^{(q)}}$, where $N_p^{(q)}$ is the total number of samples within the population $p$ that includes sample $q$; note that with this weighting $\sum_p N_p = N$ and $\sum_k w_k = 1$. In this case, Equation 29 above can be re-written as:

$$\varrho_{ij}^{(q)} = PN_p^{(q)}(\varrho_{ij}^{(\alpha)} - \varrho_{ij}^{(\alpha-q)}) + \varrho_{ij}^{(\alpha-q)}. \tag{30}$$

This approach assigns higher $w_q$ values to samples from smaller subpopulations and lower $w_q$ values to samples from larger subpopulations when calculating the weighted Pearson correlation. The consequence for LIONESS::WPCC (Equation 30) is that the differential correlation term is multiplied by a smaller value when computing the network for a sample coming from a smaller subpopulation and multiplied by a larger value when computing the network for a sample coming from the larger subpopulation. Although initially this scaling of the differential-correlation term in the context of different subpopulation sizes seems similar to the role of the $S_q$ parameter in SWEET, the ultimate consequence on the predicted network is not comparable. This is because SWEET uses a standard (unweighted) Pearson correlation to compute the aggregate and background networks while LIONESS::WPCC uses a weighted Pearson correlation to compute the aggregate and background networks. Thus the differential and background correlation terms in LIONESS::WPCC and SWEET are not the same.

### 4 Estimating single-sample networks based on measures of non-linear correlation

#### 4.1 LIONESS::MI

As noted above, LIONESS was originally derived in a method-agnostic manner. In this paper, we apply LIONESS to networks derived using Pearson correlation (LIONESS::PCC, see Section 2) as well as to networks computed using MI (referred to as LIONESS::MI). When applying LIONESS to MI networks, the following variable substitutions can be made: (1) $e_{ij}^{(q)} \sim I_{ij}^{(q)}$, the estimated sample-specific network, (2) $e_{ij}^{(\alpha)} \sim I_{ij}^{(\alpha)}$, the MI between two entities (e.g. genes), $i$ and $j$, across all samples, and (3) $e_{ij}^{(\alpha-q)} \sim I_{ij}^{(\alpha-q)}$, the MI between two entities, $i$ and $j$, across all samples except for $q$. MI gives each sample equal weight, so $w_q = \frac{1}{N}$. Thus, in the context of MI, Equation 1 can be recast as:

$$I_{ij}^{(q)} = N(I_{ij}^{(\alpha)} - I_{ij}^{(\alpha-q)}) + I_{ij}^{(\alpha-q)} \tag{31}$$

In implementing LIONESS::MI we used code from `https://github.com/otoolej/mutual_info_kNN` to calculate MI (i.e. $I_{ij}^{(\alpha)}$ and $I_{ij}^{(\alpha-q)}$) and applied Equation 31 as written.

To illustrate the mathematical synergy between LIONESS::MI and CSN (see below), we note here that in Supplemental Equation E48 of their paper, Kuijjer et al. provide an equation for LIONESS when applying it to MI estimated using a discrete binning scheme:

$$I(q) = log\left(\frac{A_{xy}^{(q)}N}{X_y^{(q)}Y_x(q)}\right) + \sum_{i\neq q} log\left(\frac{N}{N-1}\frac{A_{xy}^{(i)}}{A'^{(i)}_{xy}}\frac{X'^{(i)}_{y}}{X_y^{(i)}}\frac{Y'^{(i)}_{x}}{Y_x^{(i)}}\right) \tag{32}$$

Recasting this using variables that reflect an edge between genes $i$ and $j$ for sample $q$, Equation 32 can be re-written as:

$$\begin{aligned} I_{ij}^{(q)} &= log\left(\frac{Nn_{ij}^{(q)}}{n_i^{(q)}n_j^{(q)}}\right) + \sum_{k\neq q} log\left(\frac{N}{N-1}\frac{n_{ij}^{(k)}}{n'^{(k)}_{ij}}\frac{n'^{(k)}_{i}}{n_i^{(k)}}\frac{n'^{(k)}_{j}}{n_j^{(k)}}\right) \\ &= log(Nn_{ij}^{(q)}) - log(n_i^{(q)}n_j^{(q)}) + \sum_{k\neq q} log\left(\frac{N}{N-1}\frac{n_{ij}^{(k)}}{n'^{(k)}_{ij}}\frac{n'^{(k)}_{i}}{n_i^{(k)}}\frac{n'^{(k)}_{j}}{n_j^{(k)}}\right) \end{aligned} \tag{33}$$

where $n_{ij}^{(k)}$ is the total number of samples in the box that includes sample $k$ defined based on the bins for the expression of gene $i$ and gene $j$, $n_i^{(k)}$ is the total number of samples within the bin for gene $i$ that includes sample $k$, and $n_j^{(k)}$ is the total number of samples within the bin for gene $j$ that includes sample $k$; $n'^{(k)}_{ij}$, $n'^{(k)}_{i}$, and $n'^{(k)}_{j}$ are defined equivalently except in the context of removing a single sample ($q$) from the dataset. Some samples are in the same box or bin as $q$ while others are not. In other words, $n'^{(k)}_{ij} = n_{ij}^{(k)} - 1$ for samples in the same box as sample $q$ but $n'^{(k)}_{ij} = n_{ij}^{(k)}$ otherwise; $n'^{(k)}_{i} = n_i^{(k)} - 1$ for samples in the same bin for gene $i$ as sample $q$ but $n'^{(k)}_{i} = n_i^{(k)}$ otherwise; and $n'^{(k)}_{j} = n_j^{(k)} - 1$ for samples in the same bin for gene $j$ as sample $q$ but $n'^{(k)}_{j} = n_j^{(k)}$ otherwise.

#### 4.2 CSN

CSN (cell-specific network) was developed with the goal of inferring networks for each individual cell in a single-cell gene expression dataset. Equation 5 in Dai et al. [6] defines the network for cell $k$ as:

$$\hat{\rho}_{xy}^{(k)} = \frac{\sqrt{n-1}(n \cdot n_{xy}^{(k)} - n_x^{(k)} \cdot n_y^{(k)})}{\sqrt{n_x^{(k)}n_y^{(k)}(n-n_x^{(k)})(n-n_y^{(k)})}}. \tag{34}$$

This equation represents a normalized local joint probability calculated based on the number of cells that are located within several 'bounding boxes' around cell $k$: $n$ is the total number of cells, $n_x^{(k)}$ is the number of cells within the boundaries around cell $k$ for gene $x$, $n_y^{(k)}$ is the number of cells within the boundaries around cell $k$ for gene $y$, and $n_{xy}^{(k)}$ is the number of cells within the box defined by the boundaries around cell $k$ for genes $x$ and $y$. These boundaries are programmatically determined and their size is regulated by the 'boxsize' parameter (default equal to 0.1 in Dai et al.). We note that in the implementation provided by the authors [7], CSN has a slightly different form of the equation from the one reported above. This form includes an additional small value in the denominator:

$$\hat{\rho}_{xy}^{(k)} = \frac{\sqrt{n-1}(n \cdot n_{xy}^{(k)} - n_x^{(k)} \cdot n_y^{(k)})}{\sqrt{n_x^{(k)}n_y^{(k)}(n-n_x^{(k)})(n-n_y^{(k)}) + \varepsilon}}. \tag{35}$$

In their implementation, Dai et al. set $\varepsilon = esp = 2^{-52}$, i.e. the value returned by matlab for "esp"; for consistency, we used this form of the CSN equation when implementing the method.

Although CSN was developed with the goal of inferring networks for each cell in a single-cell gene expression dataset, the mathematics of CSN can also be applied to infer sample-specific networks using bulk expression data. In this context, each 'cell' in CSN is a 'sample' in the bulk dataset. With this in mind, we can re-cast Equation 35 using the same variables as we used for LIONESS::MI (see Equation 33): $n \sim N$, $n_{xy}^{(k)} \sim n_{ij}^{(q)}$, and $n_x^{(k)} \sim n_i^{(q)}$, $n_y^{(k)} \sim n_j^{(q)}$. With these recast variables, Equation 35 can be re-written as:

$$\begin{aligned} \hat{I}_{ij}^{(q)} &= \frac{\sqrt{N-1}(Nn_{ij}^{(q)} - n_i^{(q)} n_j^{(q)})}{\sqrt{n_i^{(q)} n_j^{(q)} (N - n_i^{(q)})(N - n_j^{(q)})}} \\ &= (Nn_{ij}^{(q)} - n_i^{(q)} n_j^{(q)}) \left( \frac{N-1}{n_i^{(q)} n_j^{(q)} (N - n_i^{(q)})(N - n_j^{(q)})} \right)^{(1/2)} . \end{aligned} \tag{36}$$
