## Supplemental File 2 -- Supplemental Figures for "Challenges and Opportunities in Single-Sample Network Modeling"

Marieke Kuijjer, Margherita De Marzio, & Kimberly Glass

February 27, 2026

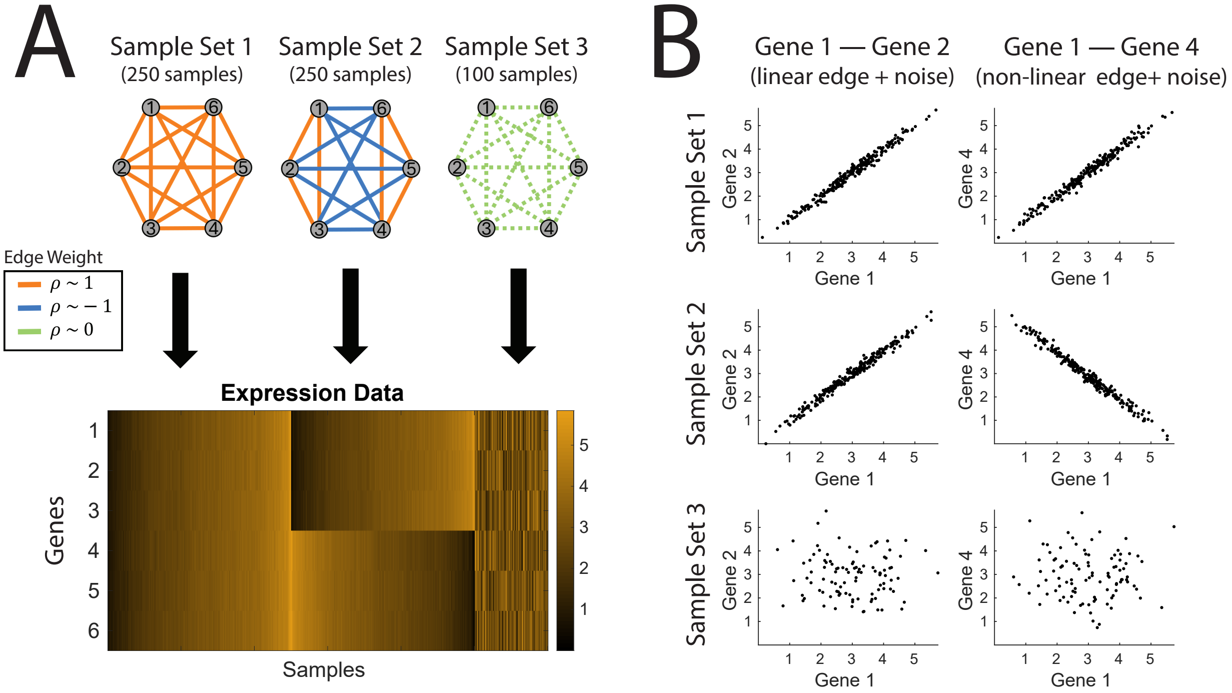


Supplemental Figure 1: **Toy Expression Data.** (A) A heat map showing the expression levels of six genes and 600 samples in the generated toy expression data. These data contain samples that represent three distinct network patterns: (1) a clique (all genes correlated with each other), (2) two cliques (two subsets of genes correlated with each other; the subsets are by definition anti-correlated), and (3) a disconnected network (genes have noisy / uncorrelated expression). (B) Examples of the two types of edges that emerge from these data, including a (1) linear edge (Gene 1 – Gene 2) in which samples from sample set 1 and sample set 2 are all highly positively correlated and samples from sample set 3 add in noise, and a (2) non-linear edge (Gene 1 – Gene 4) in which samples are positively correlated in sample set 1, anti-correlated in sample set 2, and samples from sample set 3 add in noise.

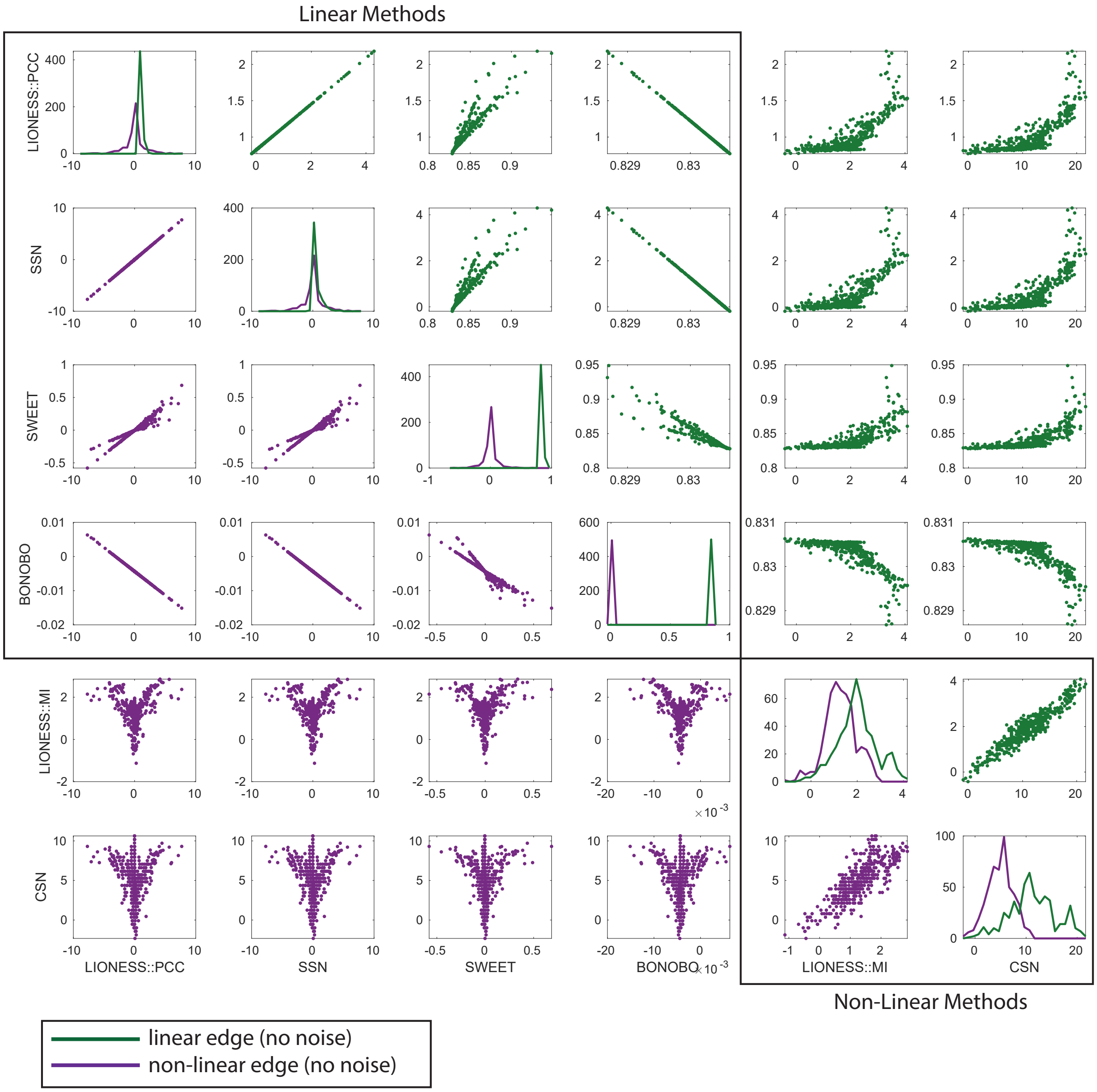


Supplemental Figure 2: Direct pairwise comparison of the edge weights predicted using each of the six single-sample network methods. Comparisons for the linear edge are shown in green in the upper-right triangle. Comparisons for the non-linear edge are shown in purple in the lower-left triangle. For each method, the distribution of the predicted edge weights across samples for each edge type are shown along the diagonal. For clarity of presentation, samples in sample set 3 (noise) are excluded from the plots.

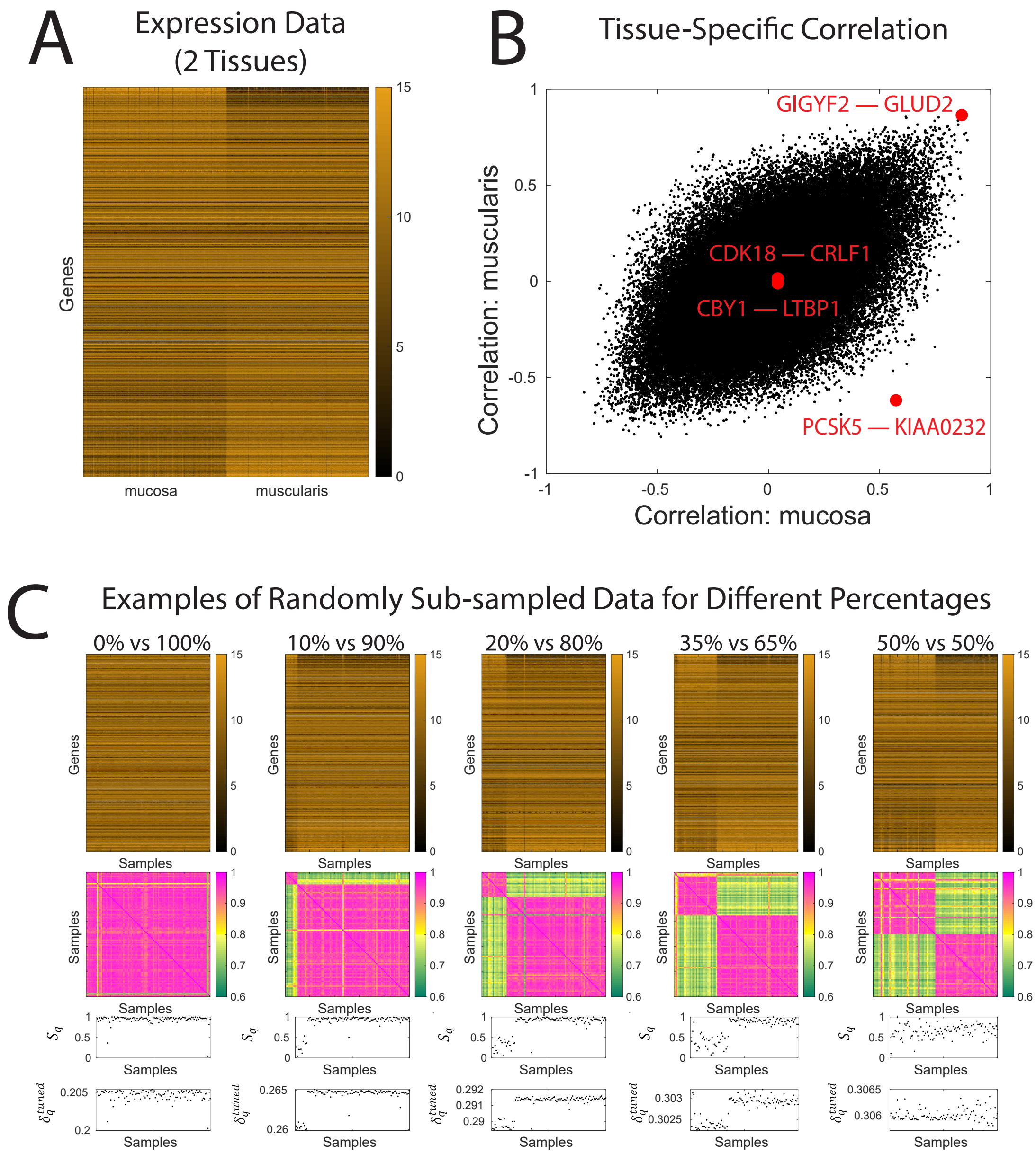


Supplemental Figure 3: (A) A 'real-world' expression dataset composed of samples from two distinct but related tissues from GTEx: the esophagus mucosa and the esophagus muscularis. These data are composed of 250 samples from each tissue. (B) A random set of 100,000 edges (gene pairs), computed based on applying Pearson correlation to the data shown in A. This plot demonstrates that most edges have very similar correlation values in both tissues, although there are some differences. The four edges selected for detailed evaluation in Figure 4 are noted as red dots. (C) first row: The gene expression values for randomly generated subsets of samples which vary with respect to the percentage of samples selected from the esophagus mucosa versus esophagus muscularis. Second row: the pairwise correlation between samples in these randomly generated subsets. Third row: the value of $S_q$ across these samples. Fourth row: the value of $\delta_q^{tuned}$ across these samples. We note that in this real-world data, sometimes samples annotated as coming from a given tissue do not cluster with other samples from that tissue. This may be due to either mis-annotation or biological variability. We also point out that the color-range used for the second row differs from that used in main text Figure 3E, as samples tend to have a much higher pairwise correlation in this real-world data compared to our toy data. To highlight differential expression patterns, in (A) and the first row of (C) genes are ordered by the product of the sum and difference in mean expression across samples from the mucosa and muscularis.

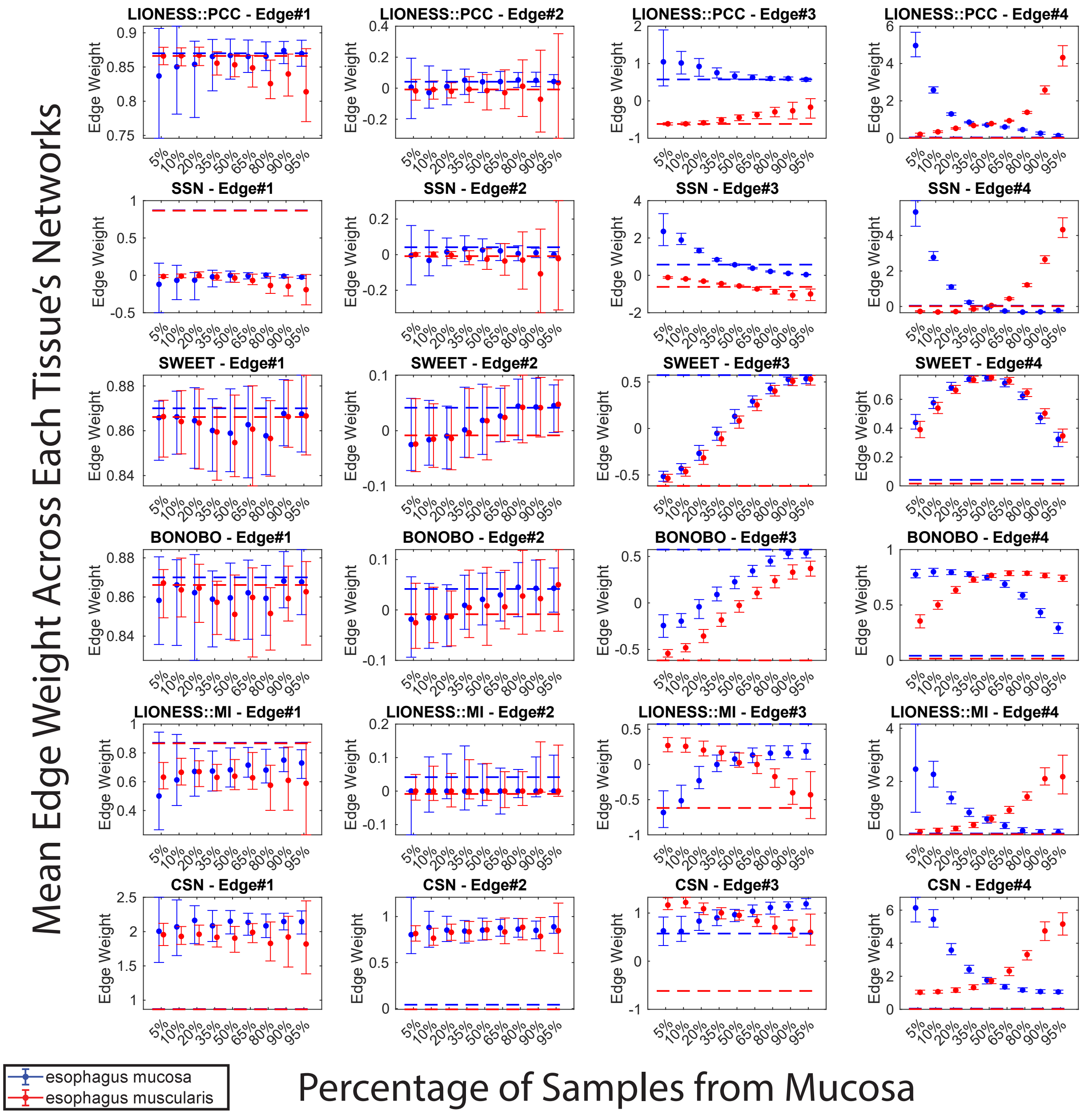


Supplemental Figure 4: Mean of edge weights across samples from each of the two tissues based on each subsampled population for the four selected exemplar edges. The median and IQR across the 100 randomizations for each percentage are shown for each method and exemplar edge. We note that the IQR calculation is sensitive to the number of samples used to calculate the underlying mean edge weights for the 100 randomizations. For example when 5% of samples are selected from the mucosa, for each randomization, the mean edge weight is calculated for only 5 samples for the muscosa but 95 samples for the muscularis, leading to a larger IQR for the mucosa versus the muscularis. The Pearson correlation (for the four linear methods) or the MI value (for the two non-linear methods) across all 250 samples from the mucosa (blue) or muscularis (red) are indicated by dashed lines. The variability in edge weight across these same sets of samples is shown in Figure 5 of the main text. The four edges correspond to the ones shown in main text Figure 5A and are as follows: Edge#1: no differential expression (DE), high correlation (corr); Edge#2: no DE, no corr; Edge#3: no DE, opposite corr; Edge#4, high DE, no corr.

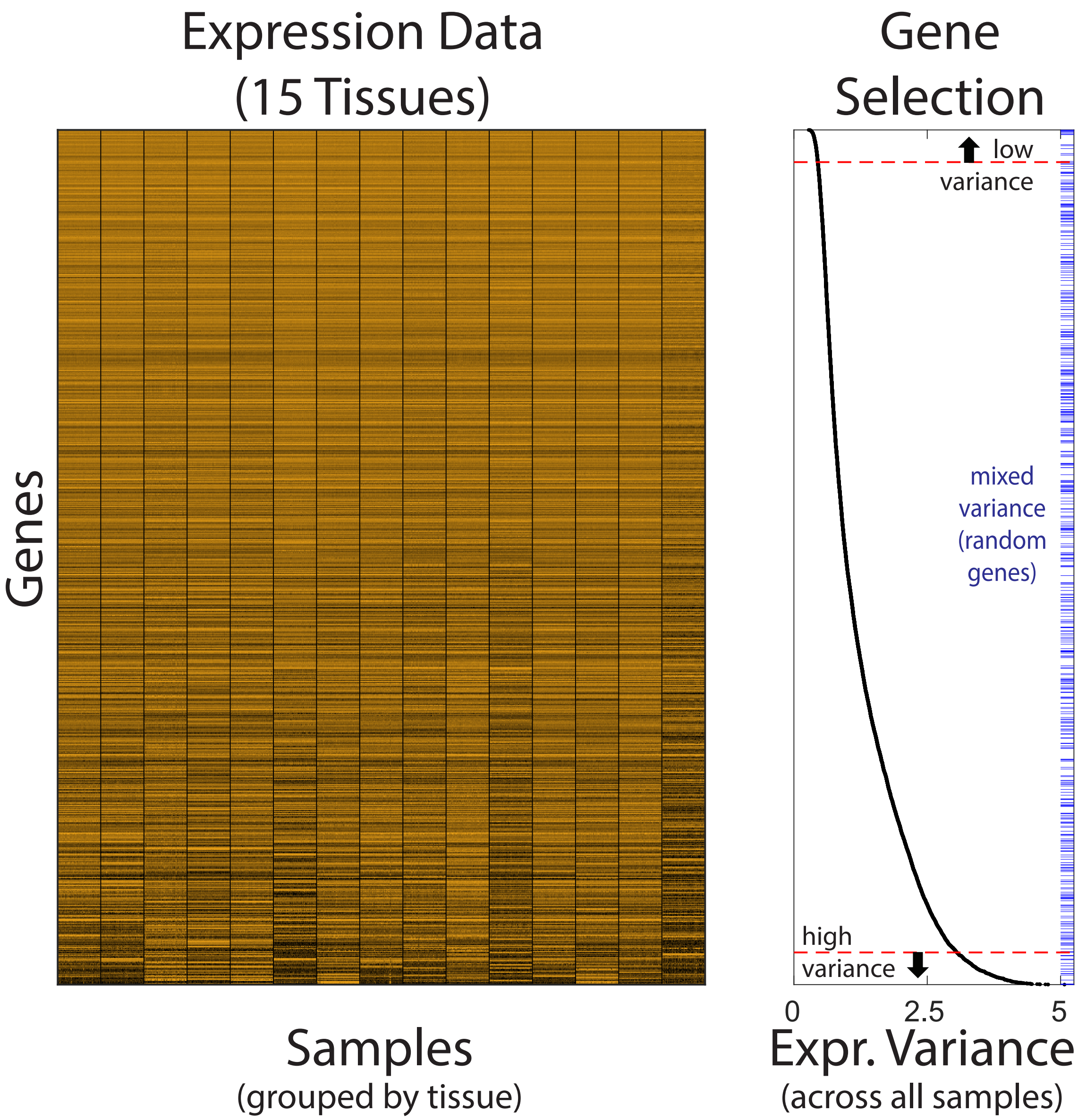


Supplemental Figure 5: A 'real-world' expression dataset composed of samples from 15 different tissues from GTEx. These data are composed of 250 samples from each tissue. Genes are ordered based on their variance across all $250 \times 15 = 3750$ samples. The groups of genes selected for detailed evaluation in Figure 5 are noted on the plot to the right, and include low variance genes (all genes above the top red dashed line), mixed variance genes (random genes, indicated by blue ticks along the right axis), and high variance genes (all genes below the bottom red line).

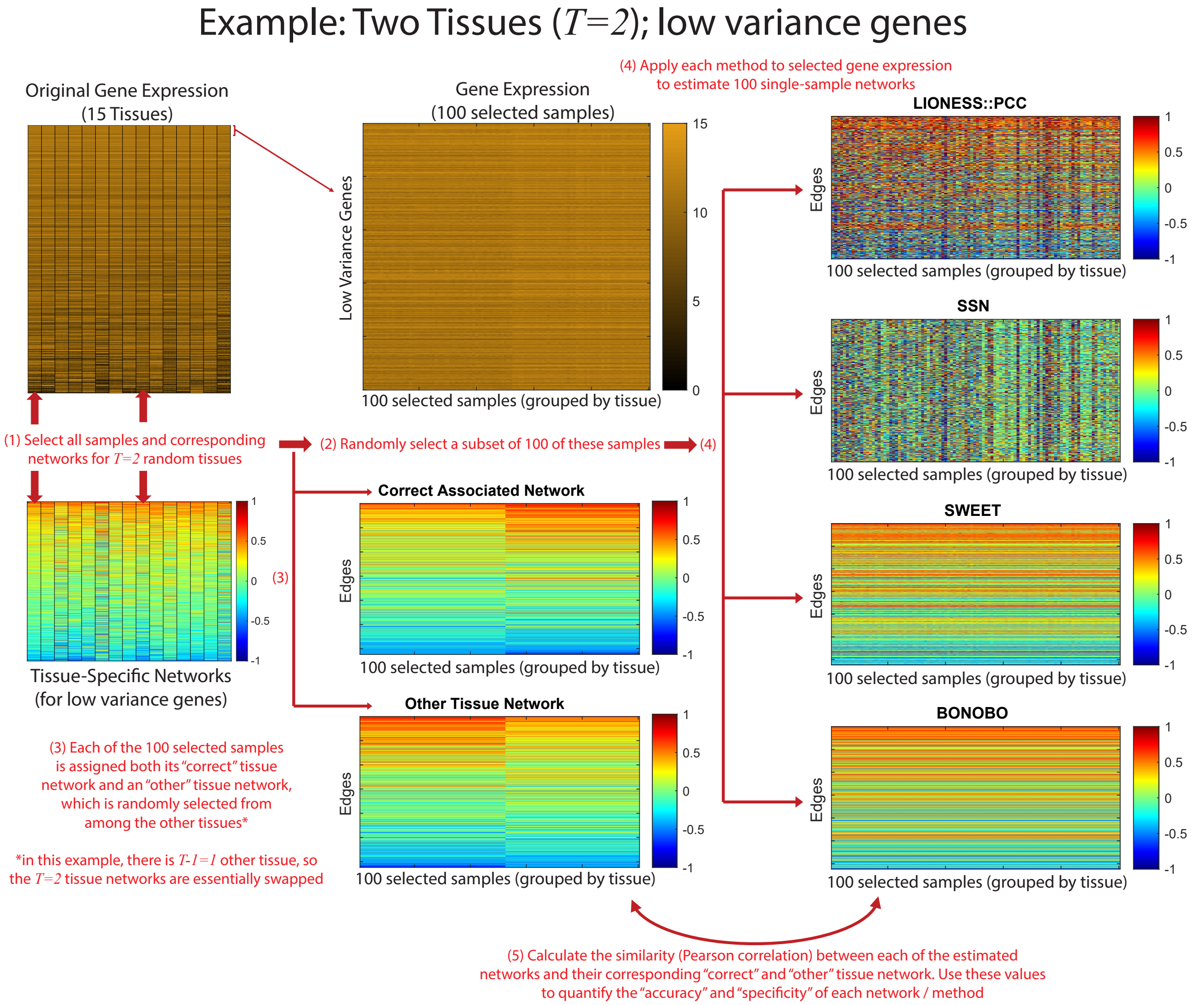


Supplemental Figure 6: Overview of the benchmarking approach taken for Figure 6 in the manuscript. An example for $T = 2$ tissues for the low-variance genes is shown.

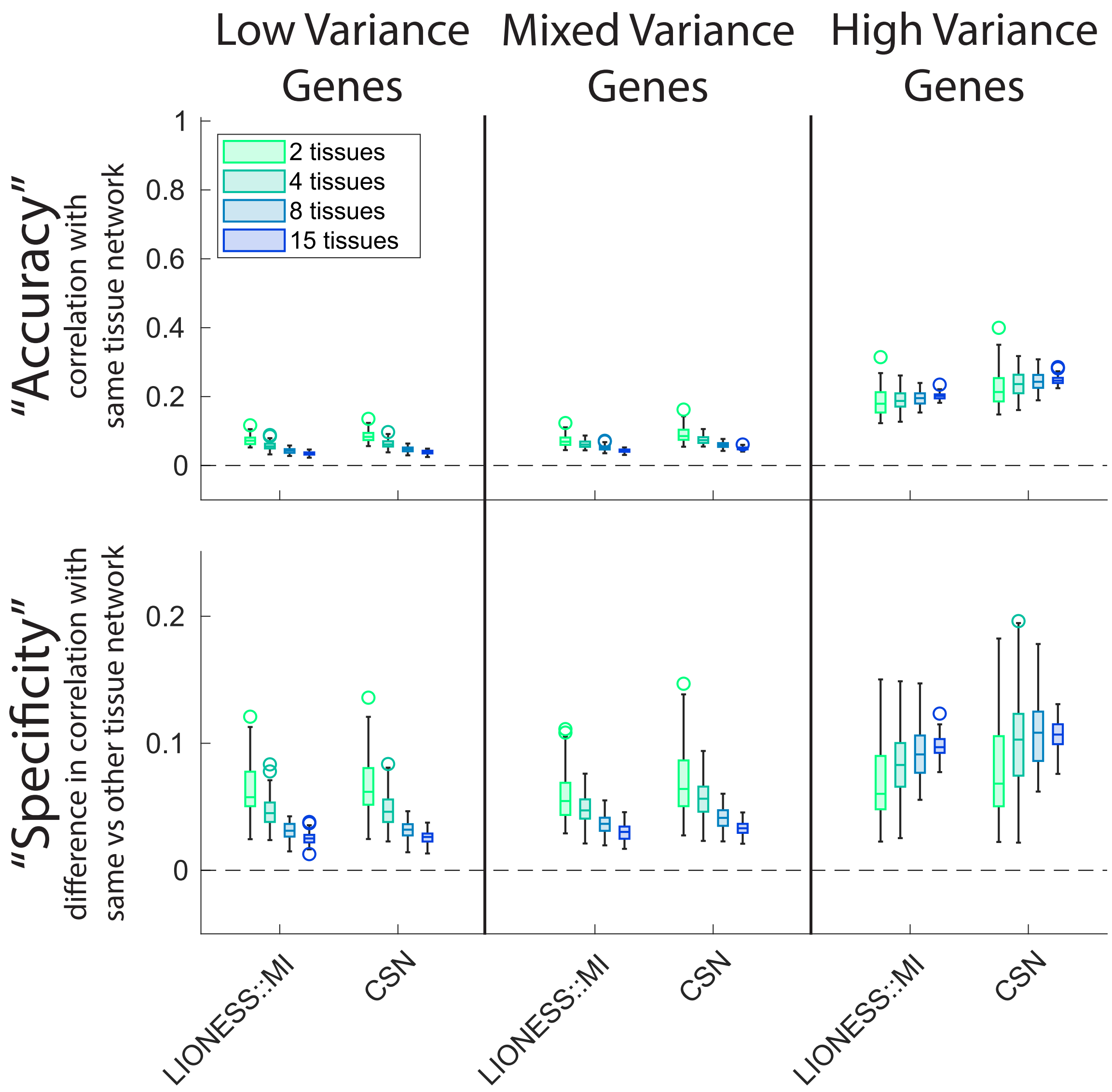


Supplemental Figure 7: The results of performing a similar benchmarking analysis as shown in the main text Figure 6 but for the non-linear single-sample network methods. In this case, the reference tissue-specific networks were MI networks (rather than Pearson correlation networks). In addition, because the non-linear single-sample network methods are more computationally intensive, only 50 genes were used for each gene set. In other words, the low variance genes are the 50 genes with the lowest expression variance, the mixed variance genes are 50 random genes, and the high variance genes are the 50 genes with the highest expression variance.
