## Supplemental File 3 -- Supplemental Analyses for "Challenges and Opportunities in Single-Sample Network Modeling"

### Supplemental File 3:
### Analysis of single-sample methods using different conceptualizations

Marieke Kuijjer, Margherita De Marzio, & Kimberly Glass

February 27, 2026

#### 1 Analysis of single-sample methods using an 'add-on-in' approach

The figures at the end of this document show the results of running the four Pearson-based single-sample network methods (LIONESS::PCC, SNN, SWEET, BONOBO) when conceptualized as 'add-one-in' approaches. This is the standard conceptualization for SSN; however, the mathematics of LIONESS::PCC, SWEET, and BONOBO can also be implemented based on this alternative conceptualization. In particular, for the 'add-one-in' approach, a set of reference samples is used to calculate a 'reference' network ($\rho_{ij}^{(\alpha-q)}$). Next, each sample is added iteratively to the reference samples and the combined data is used to calculate a 'perturbed' network ($\rho_{ij}^{(\alpha)}$). The mathematics of each method can then be applied to these two computed networks. Code implementing these four methods based on the 'add-one-in' conceptualization is included in our github repository at `https://github.com/kimberlyglass/single-sample-networks/`.

We ran each of the four linear single-sample network methods based on the 'add-one-in' conceptualization, with the simulated toy data as input and using each of the original sample sets in that data as the 'reference' sample set. In other words, we ran the methods three times: (1) with sample set 1 as the reference and predicting networks for samples in sample set 2 and sample set 3; (2) with sample set 2 as the reference and predicting networks for samples in sample set 1 and sample set 3; and (3) with sample set 3 as the reference and predicting networks for samples in sample set 1 and sample set 2. Figures with the results of these analyses, formatted to mirror the results of the 'leave-one-out' analysis shown in Figure 2 and Supplemental Figure 2, are included below.

We observe that the 'add-one-in' conceptualization has a significant impact on the values predicted by each of the methods, and that these values are often sensitive to patterns found in the reference samples. This is especially true for SWEET and BONOBO, which tend to return values that are close to $\rho_{ij}^{(\alpha-q)}$, i.e. the 'reference' network in this conceptualization. For SSN, when there is a pattern in the 'reference' network (i.e. when sample set 1 or sample set 2 is used as the reference), we see that the edge weights predicted for the samples not in the reference tend to become extreme, with values ranging in the 100s. For example, edge weight values are as high as 726 and as low as -660 for the non-linear edge using the 'add-one-in' approach with SSN, compared to values of $\pm 7.8$ for the standard 'leave-one-out' approach. The edge weight values predicted by LIONESS::PCC also shift in this context, but are in a similar range under both the 'add-one-in' and 'leave-one-out' approaches. Notably, in the context of a 'random' reference network (i.e. sample set 3 used as the reference), LIONESS::PCC and SSN give identical results. This makes sense based on their mathematical formulation, as in this case the values in the 'reference' network ($\rho_{ij}^{(\alpha-q)}$) will be close to zero.

#### 2 Analysis of single-sample methods using the original mathematics

For both BONOBO and SWEET, we slightly adjusted their original mathematical formulation to better emphasize their similarities with other single-sample network methods. For BONOBO, in the main text we presented the BONOBO equation as it would be written in the limit of large $N$ (a large number of samples). For SWEET this involved not re-replicating a sample of interest when computing a 'perturbed' network, but instead using the standard 'leave-on-out' approach. For more information, see the detailed description of how we re-cast these methods in Supplemental File 1. To ensure that these small shifts in the mathematical formulation of BONOBO and SWEET did not impact our main text results, we re-ran the entire evaluation pipeline using code for these methods that implemented their original mathematical formulation. The results are visually identical, meaning that there is absolutely no impact on the conclusions we present in our main text. We have supplied the code and figures from this analysis on a github here: `https://github.com/kimberlyglass/single-sample-networks/tree/main/Supplemental`.

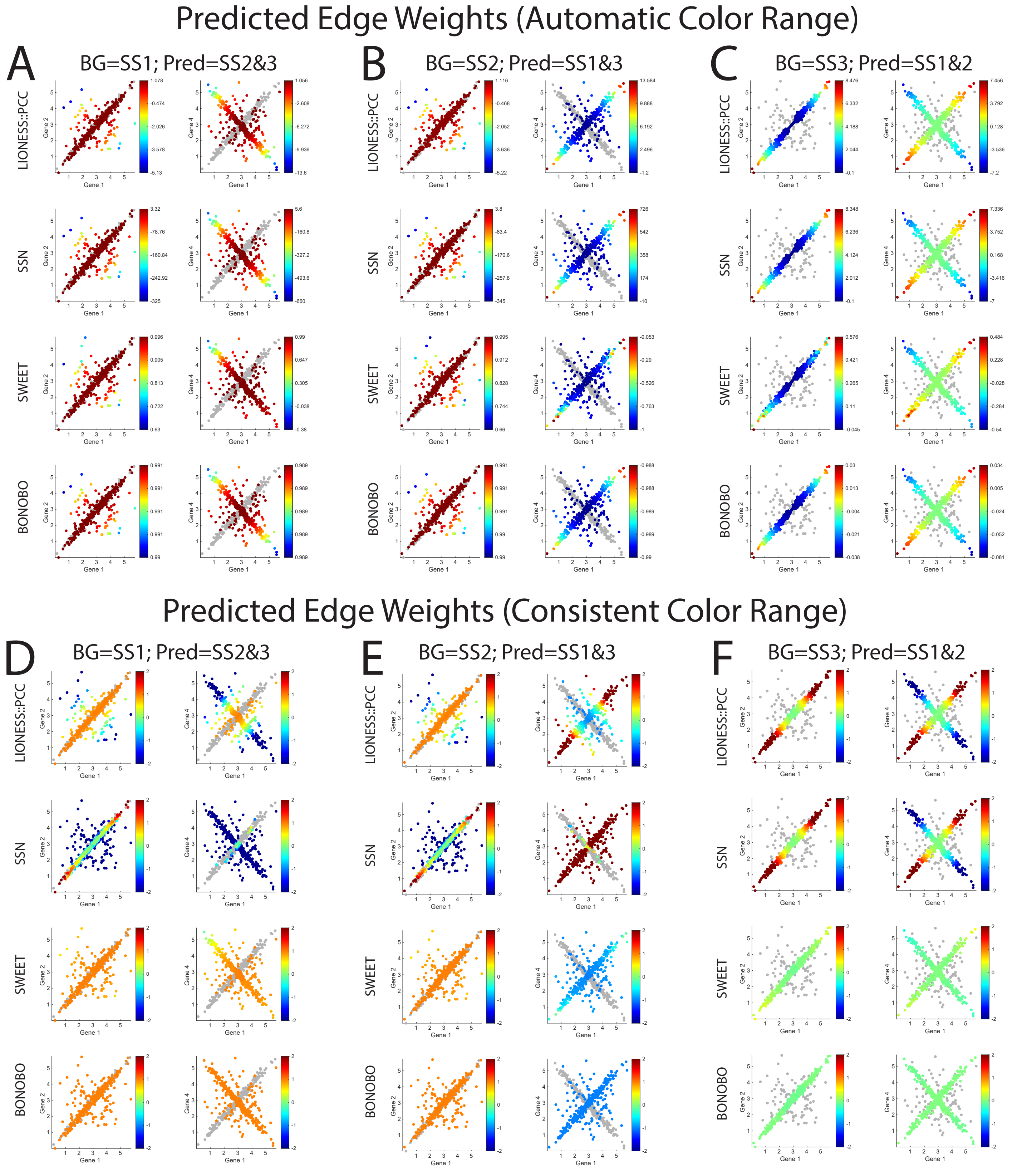


Figure 1: A visualization of the edge weights predicted for all edges, samples, and methods when using an 'add-one-in' approach. The analysis was done using either the samples in sample set 1 (A), sample set 2 (B) or sample set 3 (C) as the reference (background/BG); these reference samples are shown in gray. In each case, the edge weights for all remaining samples (Pred) were inferred using each method, and colored according to their predicted edge weights. The top panels (A-C) show the results when an 'automatic' color range is used, the bottom panels (D-F) show the results when a 'consistent' color range is used. Within each panel, the linear edge plus noise (Gene 1 – Gene 2 across all samples) is shown on the left while the non-linear edge plus noise (Gene 1 – Gene 4 across all samples) is shown on the right, consistent with Figure 2A-B in the main text.

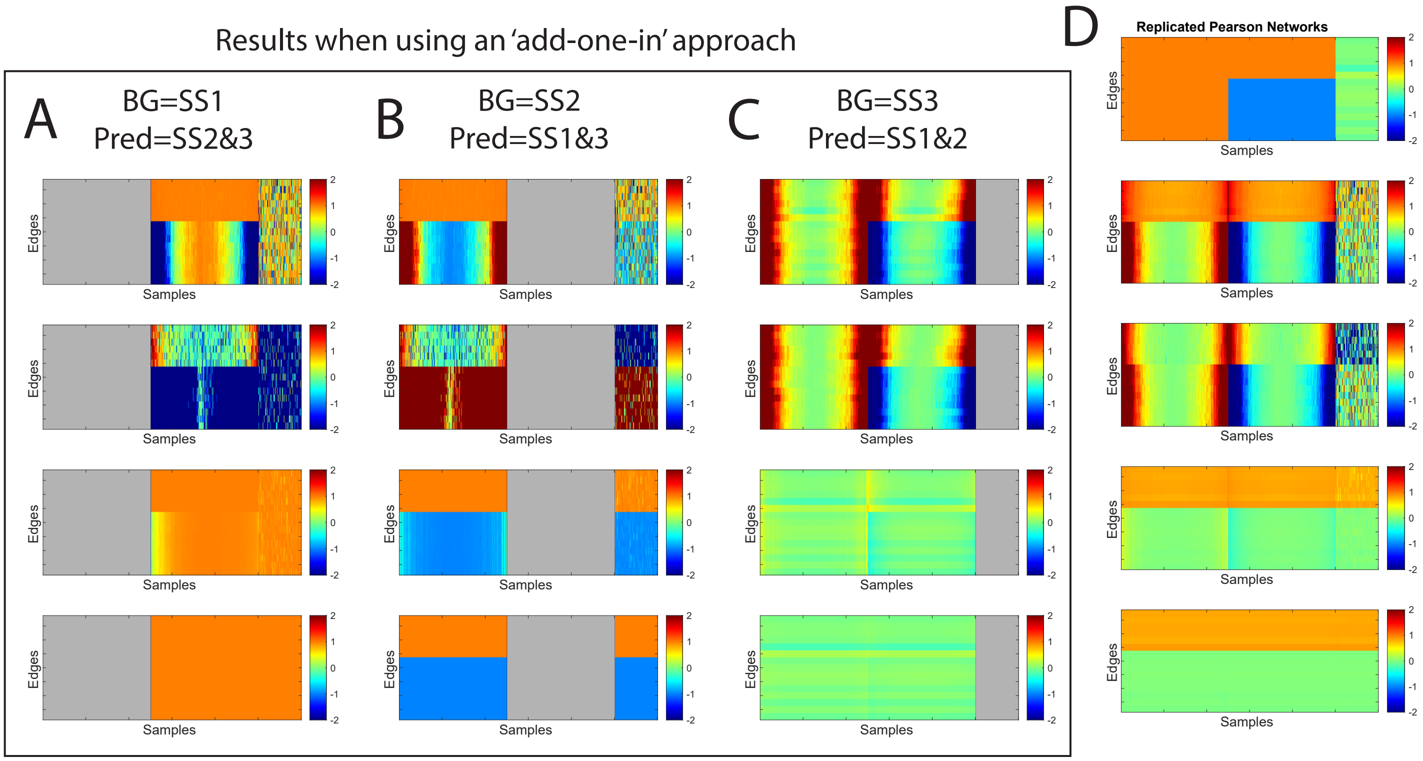


Figure 2: The edge weights predicted for all edges, samples, and methods when using an 'add-one-in' approach. The analysis was done using either the samples in sample set 1 (A), sample set 2 (B) or sample set 3 (C) as the reference (background/BG), shown in gray. In each case, the edge weights for all remaining samples (Pred) were predicted using each method. (D) A replication of Figure 2C in the main text, which visualizes the predicted edge weights for all edges, samples, and methods when using a 'leave-one-out' approach.

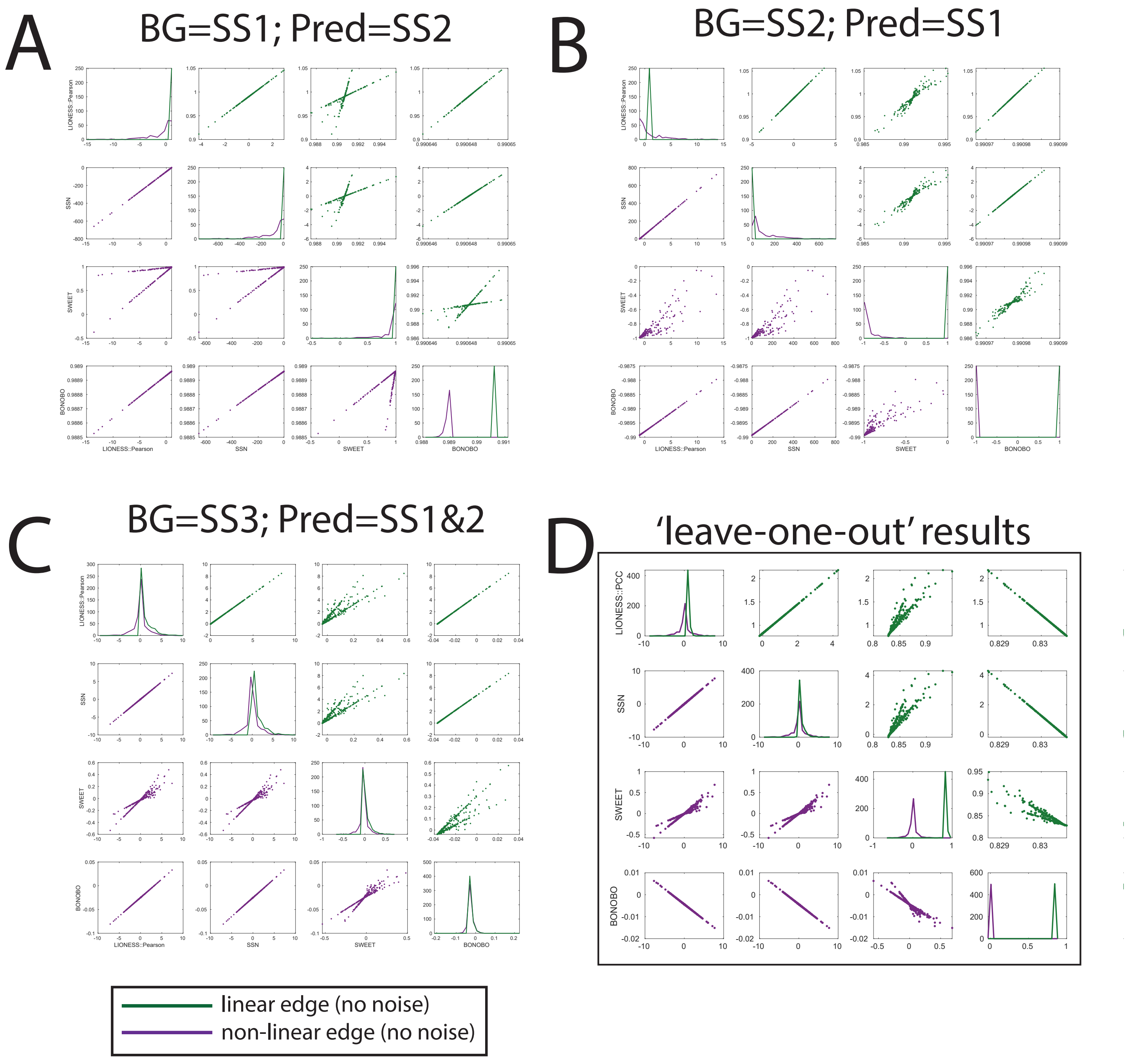


Figure 3: A comparison of the edge weights predicted for the linear and non-linear edge excluding noise (i.e. sample sets 1 and 2, but not sample set 3) when using an 'add-one-in' approach. The analysis was done using either the samples in sample set 1 (A), sample set 2 (B) or sample set 3 (C) as the reference (background/BG). In each case, the edge weights for all remaining samples (Pred) were predicted using each method, but only those belonging to sample set 1 and 2 were plotted for consistency with the primary analysis. (D) A replication of the upper left panel of Supplemental Figure 2, which compares the the predicted edge weights for the linear and non-linear edge excluding noise using a 'leave-one-out' approach.
